# Holding glycolysis in check though Alox15 activity is required for macrophage M2 commitment and function in tissue repair and anti-helminth immunity

**DOI:** 10.1101/2024.03.26.586755

**Authors:** R. Doolan, M. Moyat, G. Coakley, L. Wickramasinghe, C. Daunt, B.. Volpe, F. Henkel, V. Trefzer, N. Ubags, A. Butler, R. Chatzis, B. Marsland, A. Smith, D. Deveson Lucas, E.N.S. McGowan, K.J. Binger, J. Esser-von-Bieren, T. Bouchery, N. Harris

**Author notes:** Equal contribution (J. Esser-von-Bieren, T. Bouchery, N. Harris).

## Abstract

Macrophage polarization by type-2 cytokines is central to anti-helminth immunity and tissue repair. While some hallmark changes in macrophages are well-characterized and associated with protection against helminths, it is still unclear how macrophages exert their anti-helminth effects. In this context, we investigated Arachidonate 15-lipoxygenase (Alox15), a lipoxygenase well known for its role in macrophage polarization in the context of metabolic diseases, and a hallmark of type-2 macrophage (M2) human polarization. We show that in the absence of Alox15, M2 cannot trap and kill helminths. Surprisingly, expression of M2 markers was normal despite a loss of function. Instead, we found a concomitant increase in pro-inflammatory responses due to an uncontrolled activation of glycolysis. We further show that activation of Peroxisome proliferator-activated receptor-delta (PPAR-δ) by lipids downstream of Docosapentaenoic acid (DPA) can restore normal glycolysis control, highlighting a novel role for lipids in the fine-tuning of the metabolic support required for optimal macrophage polarization.

## Intro

Helminths are large multicellular organisms that have co-evolved with their host displaying very complex host-pathogen relationships. Because of this, despite rarely causing mortality, helminths cause a high morbidity level, especially in tropical and subtropical areas with lower economic status, where burdens are still high [1]. Despite intensive research, to date, there is no vaccine for helminth infections of humans and our understanding of protective responses against re-infection are still incomplete. Laboratory models of the three main type of helminthiases (soil transmitted helminth (STH), filariasis and schistosomiasis) have shown that communication between the adaptive and innate immune systems is required for such protection to occur. Macrophages, in particular, have been shown to be central to protective immune responses against helminths in various murine models, such as filariae and STH [2, 3]. Alternative or “M2” activation triggered by the type 2 cytokines IL-4 and IL-13, by the parasite itself, or by specific antibodies, have been shown to be required to control parasite motility and survival [4–6]. These M2 macrophages express several STAT6-induced effector molecules such as Arginase 1 (Arg1), Resistin-like molecule alpha (Relm-α, Retnla) and chitinase-like protein Ym-1 (Chil3) that have been suggested to contribute to host defense against helminths. Our group and others have described a role for Arginase-1 in protection in rodent hookworm models [4–6]. In other parasite, such as Leishmania Arg1/nitric oxide synthase 2 (Nos2) balance is thought to be crucial for protection [7, 8]. Interestingly, a recent paper challenged the importance of Arginase-1 in protection against helminth [9] and this is line with our own observations that treatment blocking Arginase-1 activity was only partially reversing protection [4]. Altogether this prompted us to evaluate other key metabolic molecules in M1/M2 polarization. In an effort to make the finding more translatable (due to high controversy on the importance of Arginase-1 in human macrophages [10]), we looked for key molecules bith highly expressed in human and mouse macrophages polarized by type 2 cytokines.

In humans, the lipoxygenase Alox15 is highly expressed in response to type 2 cytokines in macrophages both *in vitro* and *in vivo* [11–18]. Alox15 is a lipoxygenase that catalyzes the stereo-specific peroxidation of polyunsaturated fatty acids (PUFAs). Alox15 functions to generate phospholipid oxidation products crucial for orchestrating the removal of apoptotic cells as well as synthesizing precursor lipids required for the production of specialized pro-resolving mediators (SPMs) that facilitate the resolution of inflammation [19]. In mice, Alox15 has been implicated in many models of metabolic syndrome such as atherosclerosis, diabetes, obesity, and hypertension [20–22] in which macrophage polarization also plays a key role. Recently, Alox15 has also been shown to be required for the maintenance of alveolar macrophages [23]. Alox15 downstream products are endogenous ligands for the transcription factors PPAR-γ and PPAR-δ [19, 24].

In the context of helminth infection, Alox15 and its products have been shown to be upregulated in lung macrophages 7 days post-infection with *N. brasiliensis* and in intestinal tissue during secondary infection with *Heligmosomoides polygyrus* [18, 25, 26]. Furthermore, lipid pathways associated with Alox15 activity have been shown to be increased in macrophages after filarial infection [27]. However, the role of Alox15 in M2 polarization and in protection against helminth has never been addressed to date. In the current study, we demonstrate the crucial role of the Alox15 pathway in M2 polarization through control of a glycolysis:oxidative phosphorylation (OXPHOS) balance and its associated protective role in the context of parasitic infection.

## Results

### Alox15 expression is required for protection against hookworm re-infection

*N. brasiliensis* (Nb) is a rodent hookworm model that causes long-term activation of IL-4 polarised macrophages in the lungs, through which the parasite migrates on its way to the intestine. Alox15 upregulation in the lungs of Nb infected mice has previously been reported at day 7 (D7) post-infection [25]. We confirmed this upregulation of Alox15 in lung tissue and saw a low but persistent expression of Alox15 transcripts up to D40 post-infection and 2 days post re-infection (Figure S1a). Because Alox15 can be expressed by many cell types, we performed single cell sequencing of CD45+ cells to determine which cell type was principally expressing Alox15 after Nb infection and reinfection (Figure 1a&b). Consistent with the literature, macrophages and T cells are the main cell type that increase in number 40 days post-infection and 2 days post-reinfection. Alox15 expression was specifically increased upon infection in interstitial macrophages (IM), dendritic cells (DC type 1), granulocytes as well as natural killer (NK) cells, with the highest expression in IM. We further validated this increase in macrophage-specific expression of Alox15 by qPCR (Figure S1a).

**Figure 1:**
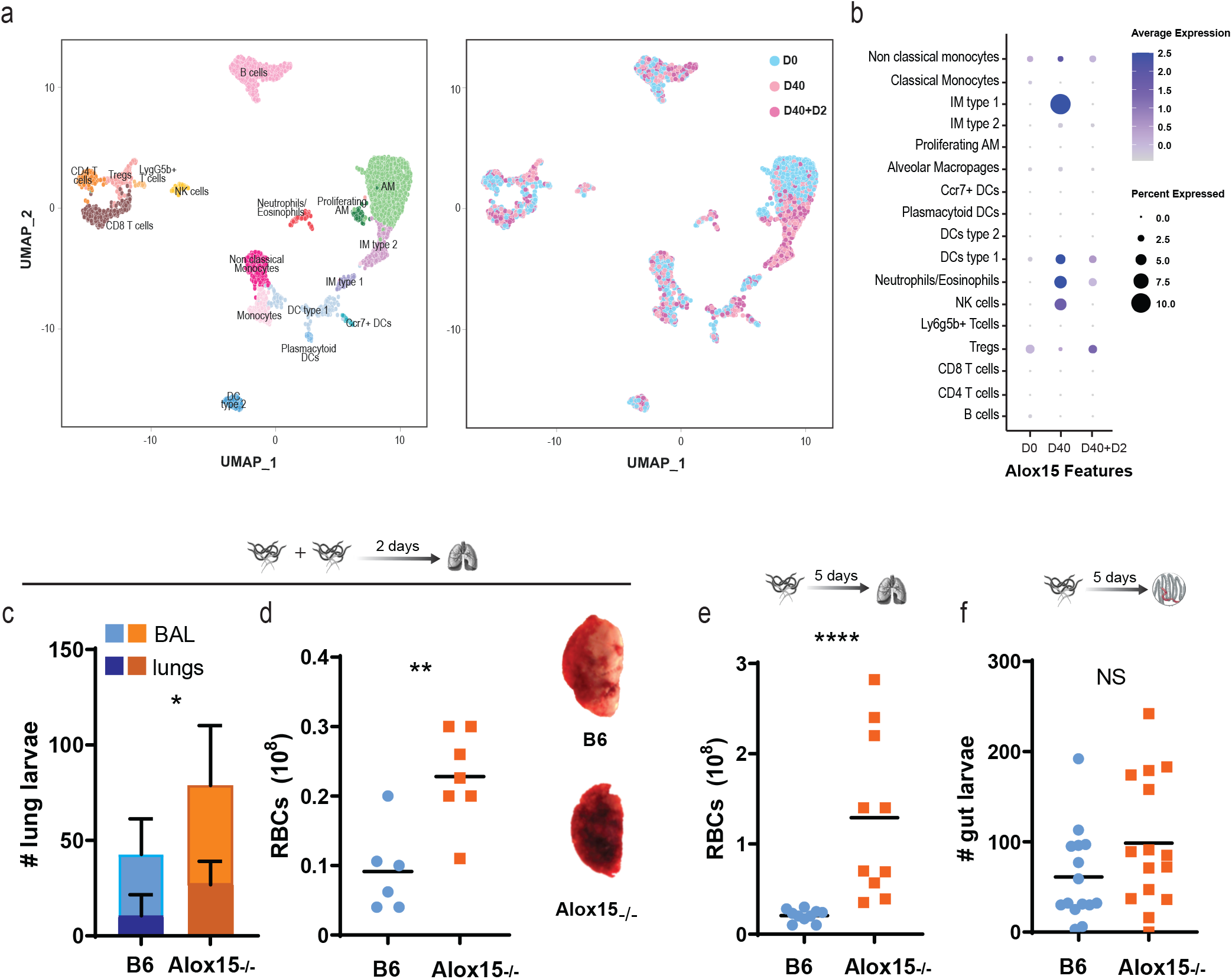
Alox15 deficiency leads to impaired protection against rodent hookworm, and reduced wound healing in the mouse lung. **A.** B6 mice were infected with 550 Nb L3s subcutaneously of left untreated (D0). 40 days post infection, mice were either left untreated (D40) or reinfected subcutaneously with 550 Nb L3s (D40 +D2) for 2 days. Lungs were harvested and CD45+ cells were isolated and prepared for single-cell sequencing on the BD Rhapsody platform (n= 4 mice per condition). UMAP visualization of scRNA-seq data of D0 (n=XXX), D40 (n=XXX) and D40+D2 (n=XXX) identifying 20 distinct cell clusters or visualized by treatment conditions. **B**. Normalized expression of *Alox15* transcripts across all cell types and shown by treatment conditions. **C,** Alox15-/-and B6 controls were infected twice with 250 Nb L3. Two days after secondary infection, larvae were collected and enumerated from BALF and baermann of lung tissue. N=5 mice per group, 2 independent experiments **D.** Hemorrhage was assessed 5 days post primary infection in BALF. N=5 mice per group, 2 independent experiments. **E.** Haemorrhage was assessed 2 days after a secondary infection. Representative photographs of lungs from B6 and Alox15-/-. N=5 mice per group, 2 independent experiments **F.** Worm burden 5 days post primary infection in Alox15-/-and B6 mice. N=5 mice per group, 3 independent experiments NS, non-significant (p>0.05); *, p≤0.05; ** p≤0.05; *** p≤0.001

To explore the role of Alox15 in macrophage polarization in response to a type-2 environment, we utilized Alox15 knock-out mice (Alox15-/-) in which a neomycin resistance cassette has been inserted in exon 3 [28]. It is well-established that upon reinfection with Nb, IL-4 polarised macrophages are able to trap and kill the parasite, resulting in a decreased worm burden in the lungs. We thus assessed the consequence of Alox15 deletion on the protective response against Nb. At 2 days post-reinfection, Alox15-/-mice could not control their infection, with a higher burden of lung larvae as compared to wild-type (WT) controls (Figure 1C). During pulmonary migration, Nb causes extensive damage to the lungs that results in short-term hemorrhage. IL-4 polarized macrophages have been shown to control hemorrhage in both primary and secondary infection [29]. We thus assessed pulmonary hemorrhage and found that hemorrhage 2 days post-reinfection was higher in Alox15-/-mice than in WT counterparts (Figure 1D).

During a primary infection, inflammatory foci and elevated red blood cell levels in bronchoalveolar lavage fluid are resolved by D5. With a classical infective dose of 550 L3, Alox15-/-mice suffered a heightened level of hemorrhage from D3 post-infection, requiring us to lower the dose of infection to avoid adverse events (FigS1b). With a low dose of infection (250 L3), equivalent numbers of larvae and hemorrhage were found between WT and Alox15-/-D2 post-infection (FigS1c&d). Interestingly, the hemorrhage was not resorbed D5 post-infection in Alox15-/-mice as compared to WT counterparts (Figure 1E). This suggests that the sustained hemorrhage observed 5 days post-infection results from decreased repair rather than enhanced pulmonary damage. IL-4 polarized macrophages contribute to timely wound repair but not the trapping of lung larvae following primary infection [29]. Consistent with a role of Alox15 in IL-4-mediated polarisation of lung macrophages, the intestinal worm burden D5 post-infection was not significantly different between WT and Alox15-/-mice (Figure 1F). Altogether, this suggests that Alox15 is important for both the repair and larvae killing function of IL-4 polarised macrophages.

### Macrophage Alox15 expression is required for larvicidal activity and tissue repair

Alox15 is expressed in many cells in the lungs and the impaired tissue repair and larval trapping observed in Alox15-/-mice after primary or secondary Nb infection respectively, could be due to defects in other cell types and involve extrinsic signaling (such as modified eicosanoids in the lung micro-environment) or intrinsic signaling (due to the loss of Alox15 expression in other cells). To determine whether the loss of Alox15 in macrophages alone can recapitulate the loss of protective immunity seen in Alox15-/-mice, we used intranasal adoptive transfer of IL-4 polarised macrophages into naïve mice to mimic the protection usually observed during secondary infection [4]. Wild-type or Alox15-/-bone-marrow derived macrophages were harvested and polarized with IL-4 in vitro to generated M(IL-4) that were transferred intranasally to naïve wild-type animals, which were infected with Nb 1 day later. Mice that received wild-type M(IL-4) had lower worm burdens than those that received Alox15-/-M(IL-4) directly implicating intrinsic Alox15 signaling in the protective response of macrophages to Nb (Figure 2A).

**Figure 2:**
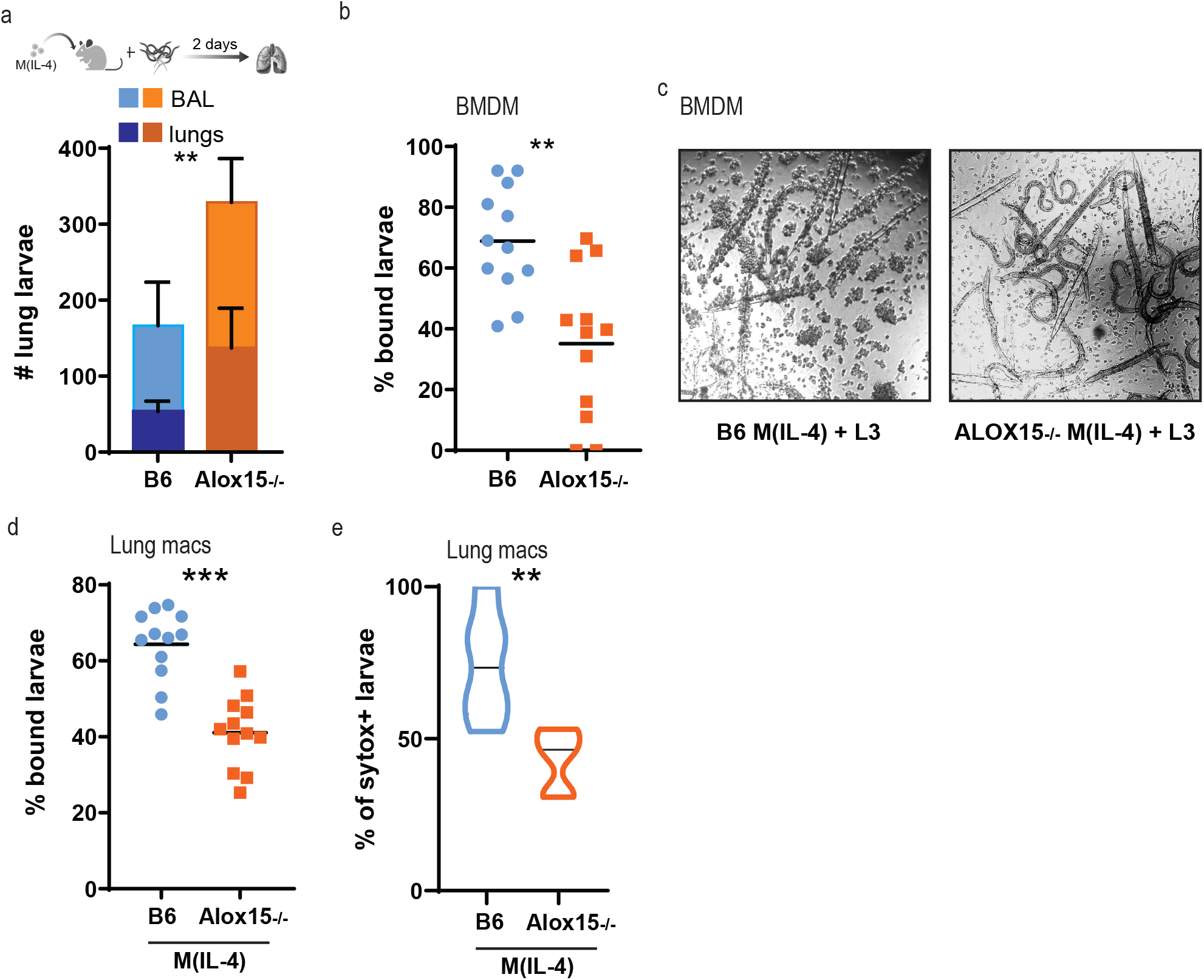
Alox15 expression in macrophages is required for larvicidal activity and tissue repair. *In vitro* and *ex vivo* co-cultures of macrophages and *Nippostrongylus brasiliensis* (Nb) larvae were used to assess the requirement of Alox15 for anti-helminth functions. Murine bone marrow-derived macrophages were grown from C57BL/6/J or ALOX-15^-/-^ mice and polarized with recombinant IL-4 for 24 hours. **A.** 2. 10^6^ BMDM (IL-4) cells were intranasally administered to naïve C57BL/6/J mice. The next day, mice were infected with 550 L3 of Nb. Two days after infection, larvae were collected and enumerated from BALF and baermann of lung tissue. N=5 mice per group, 2 independent experiments. **B.** BMDM (IL-4) were co-cultured with 100 third-stage *N. brasiliensis* larvae (Nb L3). The percentage of larvae with more than 10 cells attached was assessed for each genotypes. **C.** Representative brightfield microscopy of IL-4 stimulated BMDM (M(IL-4)) bound to Nb L3 as described in B. **D.** Lungs of uninfected B6 or Alox15^-/-^ mice were digested enzymatically and isolated macrophages used for co-culture assay as in A. (n=2-4 mice, 4 independent experiments). **E.** Viability of larvae from C., was measured using the cell impermeable fluorescent dye, Sytox Green.

Because the mechanisms by which macrophages exert larvicidal effects are incompletely understood, we further explored the importance of Alox15 in larval killing by measuring macrophage-mediated trapping *in vitro* [20]. We setup *in vitro* co-cultures of bone marrow-derived M(IL-4) with Nb third-stage larvae (L3). After 5 days of co-culture, as previously reported [20], wild-type M(IL-4) were bound to Nb L3 (Figure 2 B&C). In contrast, Nb L3 trapping by M(IL-4) generated from Alox15-/-mice was markedly reduced with less macrophages found bound to each larvae (Figure 2 B&C). As heterogeneity in macrophage populations is well established, we validated these findings by isolating pulmonary macrophages by lung digestion and examined their capacity for worm killing. As before, fewer larvae were bound by IL-4 polarized pulmonary macrophages when isolated from Alox15-/-mice compared to WT (Figure 2D). We further assessed if the larvicidal activity of Alox15-/-macrophages was compromised by measuring permeability of the larvae to the cuticle and cell impermeant SYTOX Green dye. In line with the trapping, we observed a marked increase in the percentage of dead larvae trapped by WT macrophages as compared to Alox15-/-(Figure 2E).

### Alox15 is required for larvicidal activity in an Arginase-1 independent manner

We further explored the role of Alox15 expression in regulating the phenotype of IL-4 activated macrophages. Wild-type or Alox15-/-bone marrow-derived macrophages were generated and polarized with or without IL-4. The expression of canonical M2 signature genes was studied by real-time quantitative PCR (Figure 3A). Despite a strong literature pointing towards the importance of Arginase-1 [4–6] in the anti-helminth role of macrophages, Arginase-1 expression was surprisingly higher in Alox15-/-than in WT M(IL-4) (Figure 3a). MMP-9 gene expression was significantly lower in Alox15-/-M(IL-4) than WT, while other genes typically associated with M(IL-4) polarization (Fizz1, Ym1, Fn1, MMP12, MRC1, TIMP-1) were mostly unaffected by Alox15 deletion. Due to the surprising higher gene expression of Arginase-1 in Alox15-/-M(IL-4), despite a loss of functional parasite killing, we next examined the protein expression and activity of Arginase-1. Arginase-1 expression was slightly higher in Alox15-/-M(IL-4) than WT control (Figure 3B), whilst Arginase-1 activity, measured by urea production, was similar in both WT and Alox15-/-mice (Figure 3C). To assess the relative importance of Arginase-1 and Alox15 expression in the anti-helminth function of M(IL-4), we once again set up a co-culture assay of M(IL-4) either from WT or Alox15-/-mice, with or without the addition of the Arginase-1 inhibitor BEC (*S*-(2-Boronoethyl)-l-Cysteine). As previously reported, binding of larvae by wild-type M(IL-4) was reduced by Arginase-1 inhibition (Figure 3D). However, BEC treatment did not further reduce parasite binding by Alox15-/-M(IL-4) (Figure 3D).

**Figure 3:**
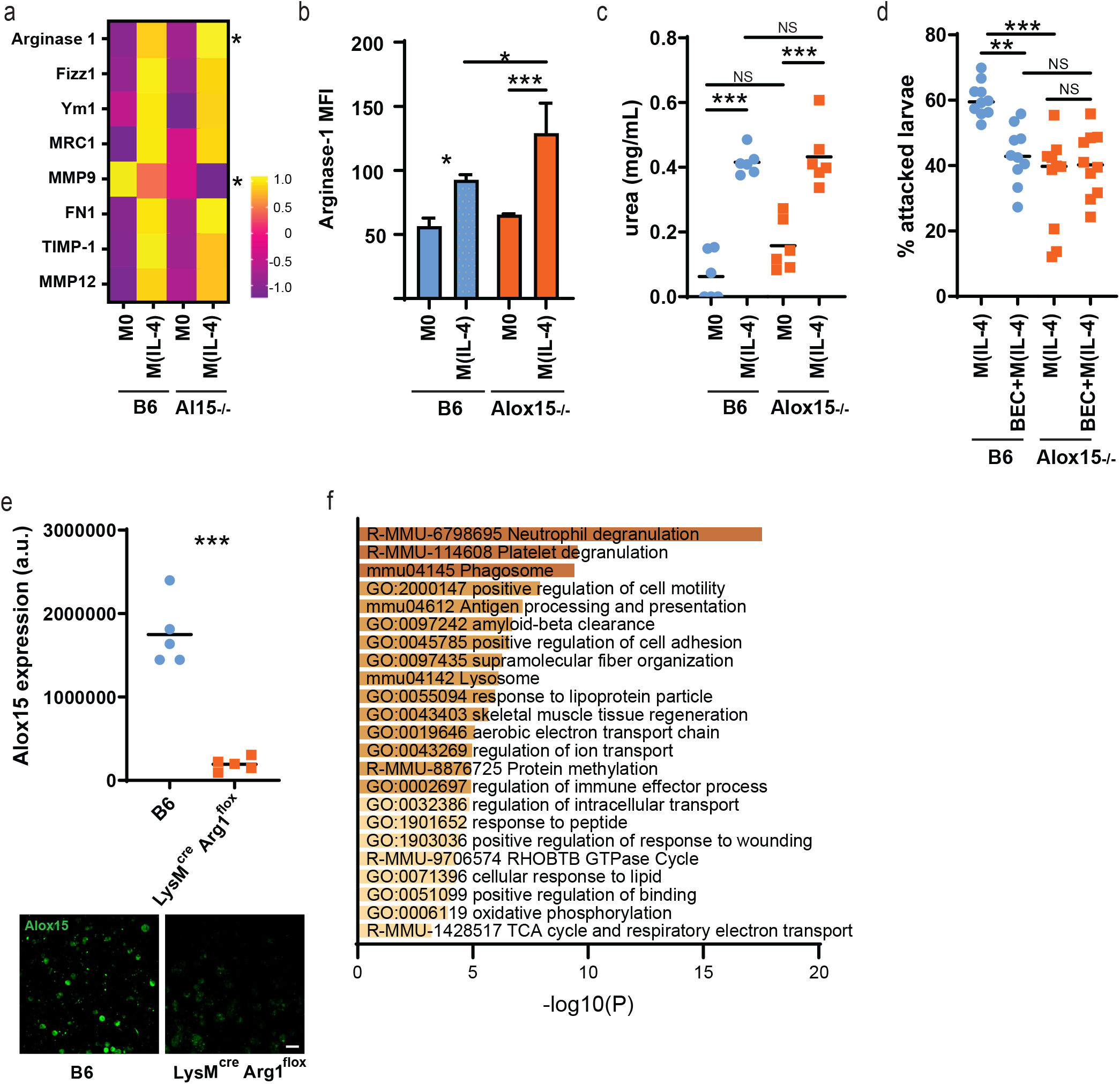
Alox15 deficiency cause a loss of function in M(IL-4) independently of Arginase-1 activity. Murine bone marrow-derived macrophages were grown from C57BL/6/J (B6) or ALOX-15^-/-^ mice and polarized or not (M0) with recombinant IL-4 (M(IL4)) for 24 hours. A. Gene expression of hallmark genes associated with M(IL-4) polarization obtained by RT-qPCR and expressed as z-score. (n=3) **B.** Mean Fluorescence Intensity of Arginase-1 analyzed by flow cytometry. (n=3) **C**. Arginase-1 activity measured by urea production. (n=3, 2 independent experiments). **D.** BMDM (IL-4) were co-cultured with 100 third-stage *N. brasiliensis* larvae (Nb L3). The percentage of larvae with more than 10 cells attached was assessed for each genotypes and conditions. (n=4-6, 2 independent experiments). **E.** Murine bone marrow-derived macrophages were grown from C57BL/6/J (B6) or LysMcre Arg1 flox mice and polarized with recombinant IL-4 (M(IL4)) for 24 hours. Immunofluorescence against Alox15 was performed and acquired on a confocal microscope. Intensity of staining was analyzed using Fiji. (n=5) **F.** Metascape pathway enrichment analysis of genes commonly upregulated in Alox15 deficient and BEC treated M(IL-4) as compared to untreated B6 M(IL-4). (n=3-4) **G.** Gene expression of hallmark genes associated with M(IL-4) polarization obtained from RNA sequencing and expressed as z-score. Note that none of those genes were identified as DEGs (n=3-4).

While Arginase-1 expression is directly triggered by IL-4 stimulation, it has previously been shown that efferocytosis is required for Alox15 gene expression to increase in response to IL-4 stimulation [19, 25]. This suggests that Arginase-1 expression could be independent, or upstream, of Alox15 expression. We thus assessed the impact of Arginase-1 blockade on Alox15 expression levels in WT M(IL-4) by immunohistochemistry. We found that Alox15 expression was reduced when Arginase-1 was absent (Figure 3E), suggesting that Arginase-1 expression is upstream of Alox15 and the role of Arginase-1 in M(IL-4) mediated trapping and killing of helminth parasites may occur through indirect pathways [5].

Despite several reports of Arginase-1 being required for parasite killing or trapping [4–6], its role in macrophage polarization following IL-4 activation has not been studied in depth. We performed bulk RNA sequencing of WT BMDM treated for 24 hours with IL-4 and/or BEC and compared it to their Alox15-/-counterparts. Surprisingly, only a few genes were significantly downregulated when comparing M(IL-4+BEC) to M(IL-4) and no pathway enrichment could be found. In contrast, 10833 were found to be upregulated by BEC treatment, although many with low fold changes (supplementary table 1). Despite BEC treatment and Alox15 depletion having similar functional consequences both *in vitro* and *in vivo* in terms of loss of parasite trapping/killing, we could not find any genes that were commonly differentially downregulated between the two treatments as compared to M(IL-4) from WT suggesting that Alox15 and Arginase-1 might fall into two distinct anti-helminth pathways or modalities, rather than Alox15 being downstream of Arginase-1.

By contrast, we found shared upregulated genes between WT M(IL-4+BEC) and Alox15-/-M(IL-4) as compared to M(IL-4) from WT and performed pathway enrichment analysis in metascape (Figure 3F). The most affected pathways were linked to degranulation, cell motility and antigen presentation, suggesting a possible role in parasite recognition. Interestingly, a few metabolic pathways (lipid metabolism, OXPHOS and TCA cycle) were also significantly upregulated in both BEC treatment and with Alox15 deficiency, suggesting that the loss of function in BEC treated or Alox15 deficient M(IL-4) could be metabolic. Of note, Alox15 expression was 1.26 times upregulated after BEC treatment as compared to WT (log fold change, p-value = 0.06) suggesting a loop of regulation between Alox15 and Arginase-1 expression. Consistent with the findings in Figure 3a, typical M2 polarisation genes were not significantly affected in Alox15-/-BMDM or in BEC treated M(IL-4) as compared to WT M(IL-4).

### Alox15 deficient macrophages have a hybrid pro- and anti-inflammatory commitment after IL-4 polarization

To refine our understanding of the role of Alox15 in IL-4 activation of macrophages, we analyzed transcriptomic changes during the transition from resting (M0) to IL-4 stimulated (M(IL-4)) BMDM for both WT and Alox15-/-mice. As expected, IL-4 stimulation caused a large number of genes to be up- and down-regulated in WT mice (Supplementary S2a & Table 2). In contrast, direct comparison of M(IL-4) from WT and Alox15-/-mice gave only 19 statistically significant differentially regulated genes (Fig S2b). Metascape was used for enrichment pathway analysis and revealed an enrichment for ‘defense response to virus’ (GO: 0051607) and ‘innate immune response’ (GO: 0045087). To refine our understanding of the role of Alox15 in macrophage polarization, we broadened our analysis to the 50 top up- and down-regulated genes between the two genotypes. Interestingly, metabolic and immune processes were found to be affected (Figure S2b). Finally, we looked at the overlap in genes affected by IL-4 treatment between WT and Alox15-/-macrophages. Both genotypes largely overlapped, with 150 shared genes but only 32 specific to Alox15 and 21 specific to WT (Supplementary S2c & Table 3). Metascape enrichment pathway analysis revealed the genes specific to WT were enriched for terms linked to cell surface interaction, potentially highlighting a role for worm recognition by macrophages. Surprisingly, genes specific to Alox15-/-were associated with ‘inflammatory response’ and ‘defense against bacteria’ (Supplementary S2c& Table 4). Potentially, such differences could explain a difference in the recognition of parasite larvae by IL-4-polarised macrophages from WT or Alox15-/-animals.

Optimal activation of ‘M2’ macrophages, in particular in the context of helminth infection, has previously been reported to require both IL-4 and/or IL-13 stimulation as well as apoptotic cell recognition [25] and is further enhanced by surfactant produced in the lungs [30]. Nb infection causes important damage to the lungs and efferocytosis has been shown to be essential in this model for optimal activation of M2 macrophages [25, 29]. Furthermore, macrophages have been shown to have tissue specificity, with lung macrophages being markedly different from bone marrow-derived macrophages [31]. To validate our findings in the context of helminth infection we assessed the role of Alox15 in IL-4 polarization *in vivo* and performed bulk RNA sequencing on lung macrophages purified 7 days post-infection with *N. brasiliensis* either in WT or in Alox15-/-mice. More marked changes were observed with 1653 genes found to be upregulated and 1073 downregulated in Alox15-/-macrophages as compared to WT counterparts (Figure 4a). We next used Metascape pathway enrichment analysis of genes specifically upregulated and downregulated in Alox15-/-lung macrophages. Upregulated genes were associated with immune response pathways characteristic of type 1 responses (response to virus, response to Interferon-beta) (Figure 4b). We thus further looked at hallmarked genes typical of ‘M2’ and ‘M1’ polarization. As reported earlier, the changes to the M2 phenotype were minor with enhancement of Arginase-1 expression and MMP-9 (figure 4c). In contrast, genes typically associated with M1 polarization were markedly higher in M2 from Alox15-/-mice as compared to WT.

**Figure 4:**
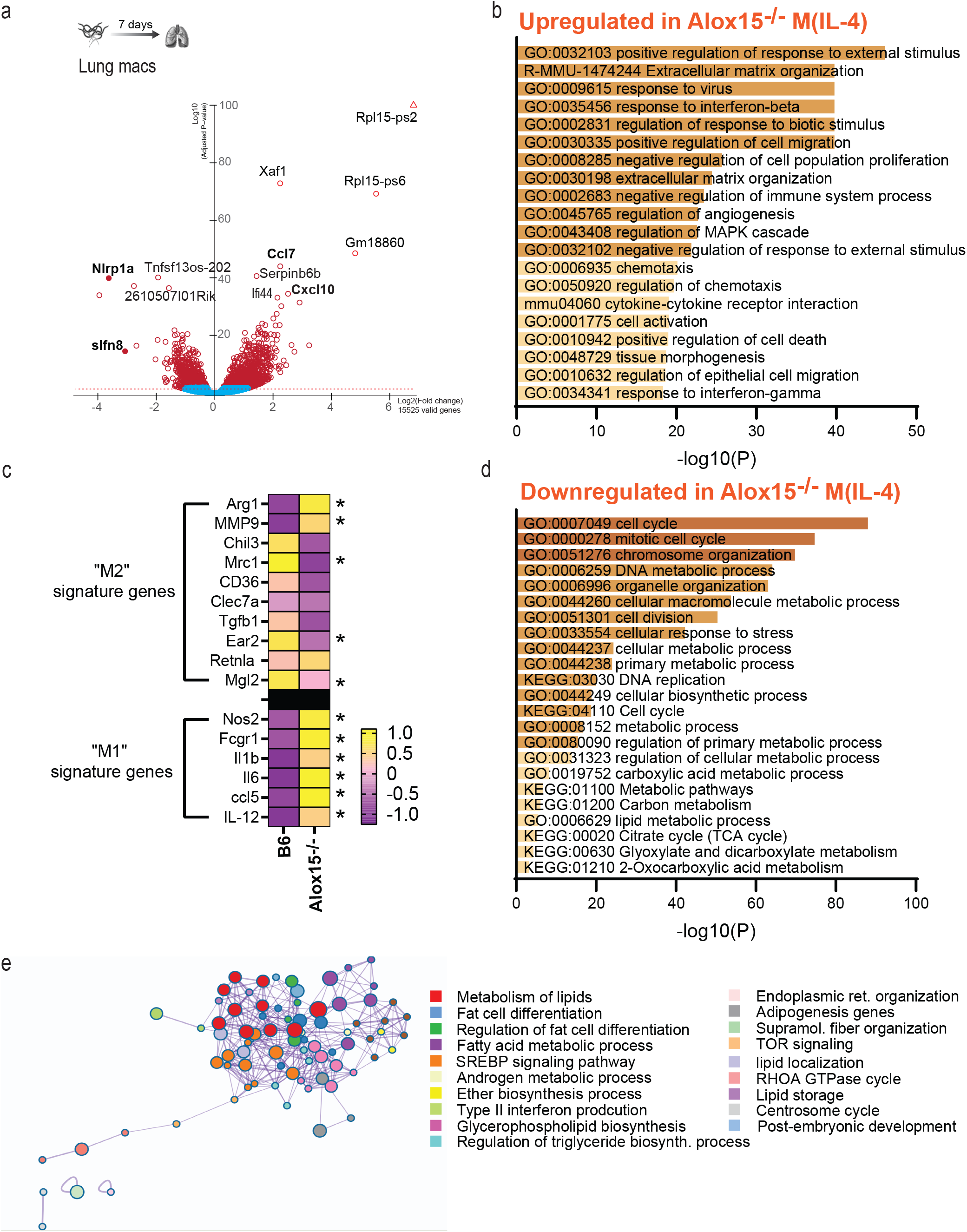
Alox15 deficiency enable development of a M1/M2 hybrid after IL-4 polarization. A. mRNA expression profiles (volcano plots) as determined by RNA-sequencing isolated from total lung macrophages from B6 or ALOX-15^-/-^ mice 7 days post-infection with Nb L3. Red, significantly differentially expressed genes with a P value <0.01 (n=4-5, representative of 2 independent experiments). B. Pathway enrichment using Metascape of DE genes upregulated in ALOX-15^-/-^ lung macrophages. C. Gene expression of hallmark genes associated with macrophages M1 or M2 polarization obtained from RNA sequencing and expressed as z-score. * corrected P-value <0.05. D. Pathway enrichment using Metascape of DE genes downregulated in ALOX-15^-/-^ lung macrophages. E. Pathway enrichment using Metascape of a cluster of DE genes downregulated in ALOX-15^-/-^ lung macrophages with a strong signature in metabolism.

As no decrease in hallmark ‘M2’ genes was observed in M2 from Alox15-/-mice, we conducted pathway enrichment analysis. The downregulated genes specifically associated with Alox15 deficiency were linked to cell cycle and cell metabolism (Figure 4d). Association between cell cycle and Alox15 deficiency in macrophages has recently been reported [23], although the authors show that at resting state this defect is limited to alveolar macrophages. Similarly, we only observed this link to cell cycle pathways *in vivo* and not in our RNA sequencing dataset derived from BMDM (Figure S2a-c). To determine whether proliferation contributed to the *in vivo* or *in vitro* phenotypes we observed, we assessed proliferation of BMDM from WT or Alox15-/-mice stimulated or not with IL-4. We observed that BMDM stimulated with IL-4 from Alox15-/-mice had no difference in Ki67 expression after one day of IL-4 stimulation. (Fig S2d). We thus concluded that a deficiency in cell cycle was unlikely to explain the lack of larvicidal activity observed in Alox15-/-M(IL-4). Furthermore, given that interstitial macrophages, rather than alveolar macrophages, are required for Nb trapping [4, 32], we decided to further focus on the metabolic changes observed in Alox15-/-lung macrophages. To separate the effect of cell cycle and metabolism we conducted cluster enrichment analysis. The genes downregulated in Alox15-/-lung macrophages formed two separate clusters enriched for cell cycle or metabolic responses respectively. Further pathway enrichment analysis on the latter cluster revealed a strong downregulation of lipid-associated pathways in Alox15-/-lung macrophages (Figure 4e).

### Alox15 deficient macrophages have a high energetic profile after IL-4 polarization

Polarization of macrophages by IL-4 or IFN-γ has been shown to induce different metabolic requirements [33]. As a main change observed by RNAseq of lung macrophages was linked to metabolic pathways, we further explored the transcriptomic signature of enzymes involved in glycolysis and OXPHOS in lung macrophages of WT or ALOX-15-/-background. In line with the increased expression of M1 polarization genes, genes associated with glycolysis were upregulated in Alox15-/-lung M(IL-4) (Figure 5a). Despite almost no change in IL-4-induced signature genes in Alox15-/-macrophages, genes associated with oxidative phosphorylation – which are typically upregulated by IL-4 – were decreased in our dataset (Figure 5b), as were genes associated with fatty acid oxidation and lipolysis (supplementary Fig S3a&b).

**Figure 5:**
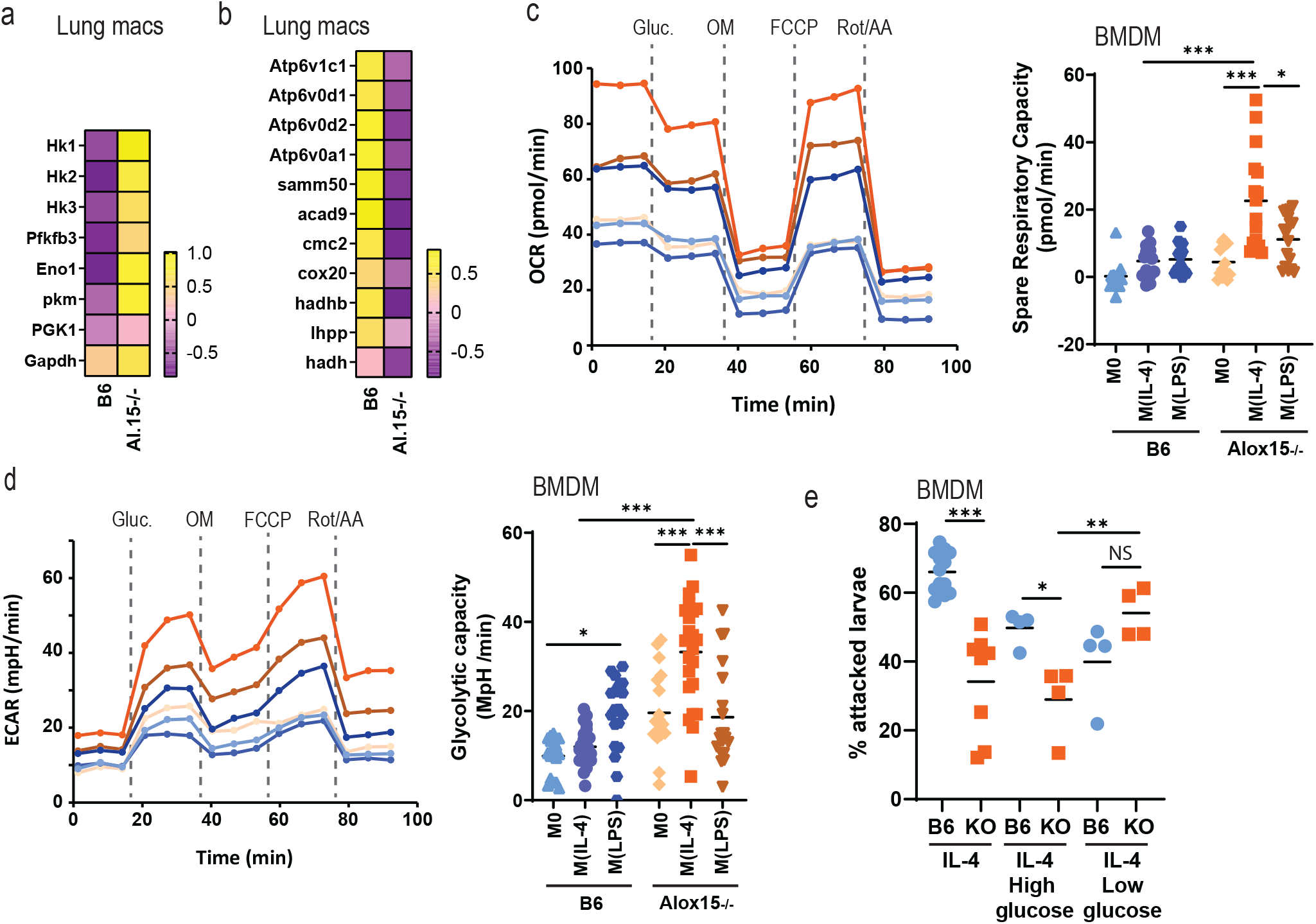
Alox15 deficient M1/M2 hybrid have a heightened energetic profile after IL-4 stimulation. Total lung macrophages from B6 or ALOX-15^-/-^ mice 7 days post-infection with Nb L3. A. Gene expression of hallmark genes associated with the glycoltic pathway obtained from RNA sequencing and expressed as z-score. B Gene expression of hallmark genes associated with OXPHOS pathway obtained from RNA sequencing and expressed as z-score. C. OCR and SRC (Spare respiratory Capacity) of BMDM stimulated or not with IL-4 or LPS from both B6 or ALOX-15^-/-^ mice at baseline and after sequential treatment (vertical lines) with glucose (Gluc.), oligomycin (OM), FCCP and Pyruvate, Antimycin-A (AA) and Rotenone (rot) as a combined Mito and Glycolytic stress test [79]. Data representative of 3-4 independent experiments, n = 5 wells per conditions, each conditions with 1-3 mice per experiments. D. ECAR and Glycolytic capacity of BMDM stimulated or not with IL-4 or LPS from both B6 and ALOX-15^-/-^ mice as described in C. E. BMDM stimulated with IL-4 in normal medium (IL-4), with IL-4 in Seahorse medium in absence of glucose (IL-4 low glucose) or in Seahorse medium with glucose (IL-4 high glucose) were co-cultured with 100 third-stage *N. brasiliensis* larvae (Nb L3). The percentage of larvae with more than 10 cells attached was assessed for each genotypes.

We next wanted to validate whether these gene expression changes translated to altered metabolic phenotypes. Using a Seahorse bioanalyser we compared resting BMDM (M0) to M(IL-4) and M(LPS) from a WT or Alox15-/-background. Even at resting state, Alox15-/-macrophages had a high energetic status with both a high ECAR and high OCR, as compared to WT (supplementary Fig S3c). This difference was even stronger after IL-4 polarization (supplementary Fig S3c). At baseline, there was no difference in spare respiratory capacity (SRC) between WT and Alox15-/-macrophages. However, after IL-4, but not LPS stimulation, there was a marked difference in SRC between the two genotypes, suggesting that Alox15 macrophages were still receptive to IL-4-mediated induction of OXPHOS as has been reported previously for WT mice (Figure 5c).

Interestingly, even from baseline, Alox15-/-BMDM exhibited a higher glycolytic capacity than their WT counterparts (Figure 5d). As expected, stimulation with LPS enhanced glycolysis in WT mice, but not in Alox15-/-macrophages (Figure 5d). Instead, a marked increase in glycolysis could be seen after IL-4 polarization in Alox15-/-macrophages, in an apparent superposition of the two metabolic responses usually attributed to M1 and M2 polarizations.

To investigate if increased glycolysis in Alox15-deficient macrophages was responsible for their lack of anti-helminth function, we polarized BMDM with IL-4 in medium containing either a lower or higher glucose content and examined worm binding as before. Interestingly, when Alox15-/-macrophages were polarized in glucose-limited conditions, cells were able to trap larvae to the same extent as their WT counterparts (Figure 5e). In contrast, WT macrophages polarized in glucose-limited conditions were less efficient at trapping larvae in line with a previous report of poor M2 polarization in low glucose settings (Figure 5e, [32]).

### Alox15 is central to the anti-helminth activity of macrophages but not in anti-*Leishmania* immunity

We found it striking that after LPS polarization, OXPHOS was heightened in Alox15-/-macrophages but that the high baseline glycolysis rate of Alox15 macrophages remained unchanged. We thus decided to further explore the importance of Alox15 in M1/M2 polarization.

First, we polarized bone marrow derived macrophages with M2 or M1 stimulants (IL-4, IL-13, LPS and IFNγ) and then co-cultured these cells with Nb. As previously described [34], ‘M0’ macrophages can to some extent bind to Nb larvae and IL-4 polarization enhances this binding (Figure 6a). Interestingly, the reduced binding of Alox15-/-macrophages was not observed when cells were stimulated with IL-13 alone or IL-13 together with IL-4, suggesting IL-13 polarisation might be Alox15-independent. In line with the literature, ‘M1’ polarized macrophages, stimulated with LPS or LPS+IFNγ, were unable to bind to Nb larvae when derived from WT mice (Figure 6a). Alox15-/-mice exhibited increased binding as compared to M1 macrophages from WT mice, and instead looked similar to M0 macrophages from either strain (Figure 6a). This indicate an overall defect in macrophage polarization in absence of Alox15.

**Figure 6:**
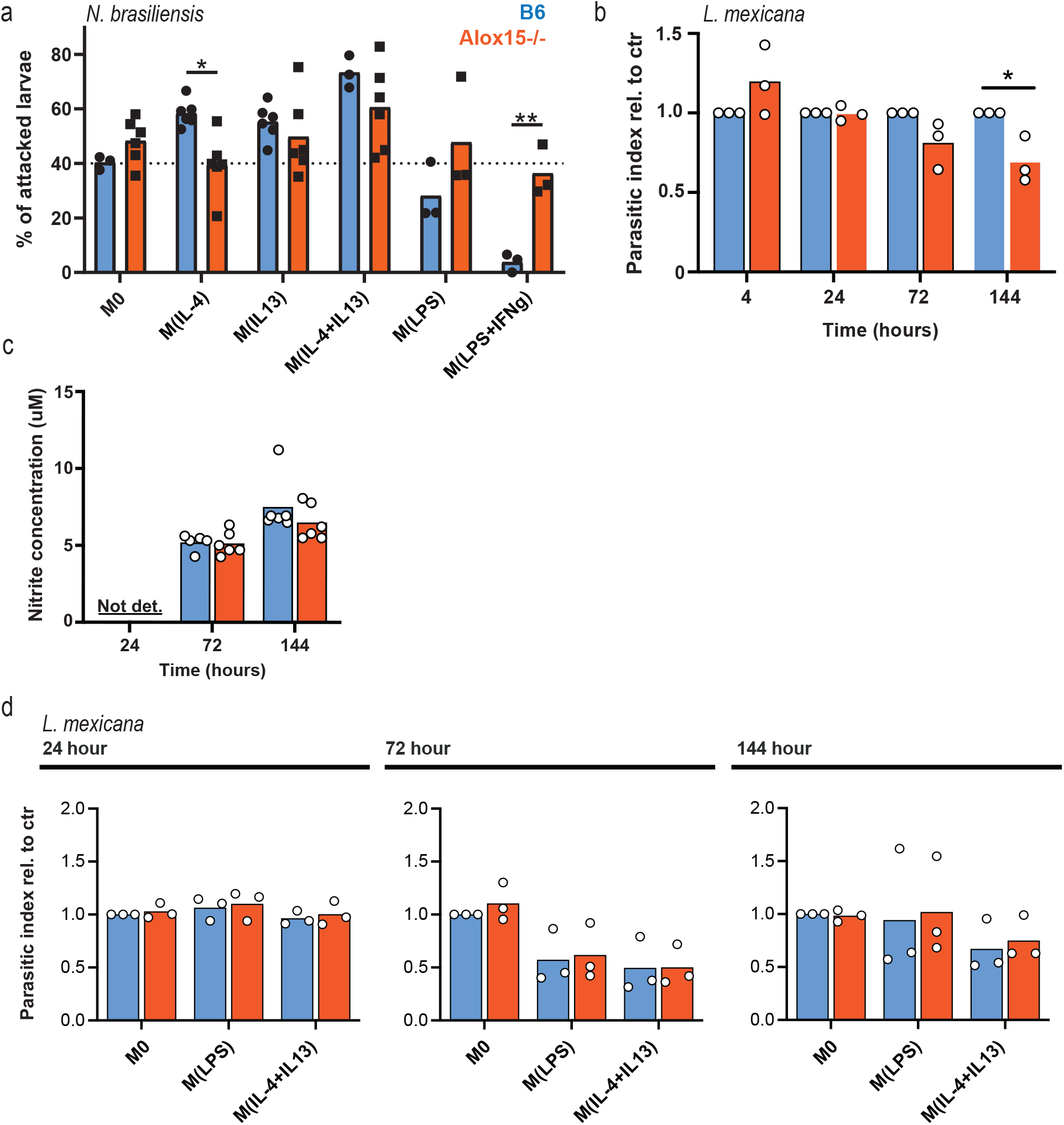
Alox15 deficiency do not impair macrophage control of *Leishmania major*. **A.** Murine bone marrow-derived macrophages were grown from C57BL/6/J (B6) or ALOX-15^-/-^ mice and polarized or not (M0) with recombinant IL-4, with IL-13, with a combination of IL-4 and IL-13, with LPS or LPS and IFNg for 24 hours. Then those were co-cultured with 100 third-stage *N. brasiliensis* larvae (Nb L3). The percentage of larvae with more than 10 cells attached was assessed for each genotypes. n=3, pool of 1-2 independent experiments. **B.** Parasitic index relative to control of a low dose of L. Mexicana infection of BMDM grown from C57BL/6/J (B6) or ALOX-15^-/-^ mice. The proportion of macrophages infected with *L. mexicana-TurboRFP* was measured 4 h to 144h later by flow cytometry. (n=3, 3 independent experiments) **C.** Nitrite concentration measure by Griess assay (n=3, 2 independent experiments) **D.** Parastic index as in B after a high dose of infection (n=3, 3 independent experiments)

Unlike helminths, Leishmania are obligate intracellular parasites. Macrophages act as the replicative niche or clear intracellular parasites, depending on their activation state. Increase ARG1 activity is thought to be beneficial to the parasite in several ways. Firstly, ARG1 competes with NOS2 for L-arginine substrate, thereby diminishing NO· and parasite death [35–37]. Secondly, increased availability of L-arginine in non-inflammatory macrophages is diverted into polyamine synthesis. Polyamines are required for parasite growth and production of the antioxidant trypanothione, which assists in withstanding the oxidative environment within the parasitophorous vacuole[35]. ARG1 catalyses the conversion of L-arginine to urea and L-ornithine, a precursor for polyamine species required for Leishmania spp. replication [35–38].

Given the mixed M1/M2 polarisation of Alox15 M(IL-4) and higher Arginase-1 expression, we set out to assess if control of this protozoan parasite is similarly affected by Alox15 deficiency. Interestingly, in a low dose model of infection with *L. mexicana*, Alox15 deficiency caused a slight decrease in parasite index at late time points indicating better macrophage control (Figure 6b). Despite this, we did not observe a difference in nitric oxide concentration even at late time points between the two genotypes (figure 6c). We next increased the multiplicity of infection to highlight differences in genotypes. Higher dose infections are more susceptible to a shift in the Arginase-1/iNOS balance. Interestingly, in high dose experiments, no difference between genotypes could be seen at any of the time points (Figure 6d). Polarization with LPS or with IL-4+IL-13 gave similar results (Figure 6d), suggesting that Alox15 is not involved in the control of Arginase-1/iNOS balance, nor is it required for killing of *L. mexicana*.

To prove that the phenotype we reported is not just specific to IL-4-mediated polarization of macrophages to a phenotype able to kill Nb larvae, we utilized another rodent helminth parasite, namely *Heligmosomoides polygyrus bakerei* (Hpb). Like Nb, protection against secondary infection requires Arginase-1 expressing macrophages [6]. As Hpb is an enteric nematode granulomas rich in macrophages are typically formed around the larvae within the submucosa of the upper small intestine. However, in contrast Nb, activation of macrophages able to kill Hpb larvae requires antibody activation, rather than IL-4. At 4 and 14 days post-secondary infection, Alox15 deficient animals had a higher worm burden that their WT counterparts (Figure 7a&c). During reinfection, granulomas rich in macrophages are typically formed around the larvae in the duodenum of the small intestine. We therefore assessed granuloma number and composition in WT and Alox15-/-mice. There was a marked delay in granuloma formation in Alox15-/-mice, with significantly fewer granulomas in Alox15-/-mice than their WT counterparts 4 days post-reinfection (Figure 7b&d). Of note, parasite burden in primary infection was not different between genotypes (Figure S4a). In line with our findings for Nb infection, the granulomas were still rich in macrophages and expressed heightened level of Arginase-1 in Alox15-/-mice (Figure 7e). As Alox15 can be expressed by other cells that could be involved in the granulomas (eosinophils and epithelial cells), we next used an *in vitro* system to assess the role of Alox15 in macrophage trapping of Hpb larvae. We again generated BMDM and co-incubated them with Hpb L3 in the presence of serum from infected mice, but without IL-4 activation. In line with the phenotype of decreased trapping with Nb, the motility of the larvae co-incubated with (immune serum activated) Alox15-/-macrophages as compared to WT was higher, suggesting a less efficient trapping (figure 7f). It has previously been shown that the decreased motility of Hpb larvae trapped by macrophages can be reversed by Arginase-1 blockade. We first assessed Arginase-1 activity after Hpb and immune serum incubation in Alox15-/-macrophages, and as in our Nb L3 and IL-4 assays, Arginase-1 activity was markedly enhanced in KO macrophages (figure 7g). We further determined that despite a notable difference in the drivers of macrophage polarization (antibodies versus IL-4), Arginase-1 blockade was not able to further increase motility in Alox15-/-macrophages (figure 7f).

**Figure 7:**
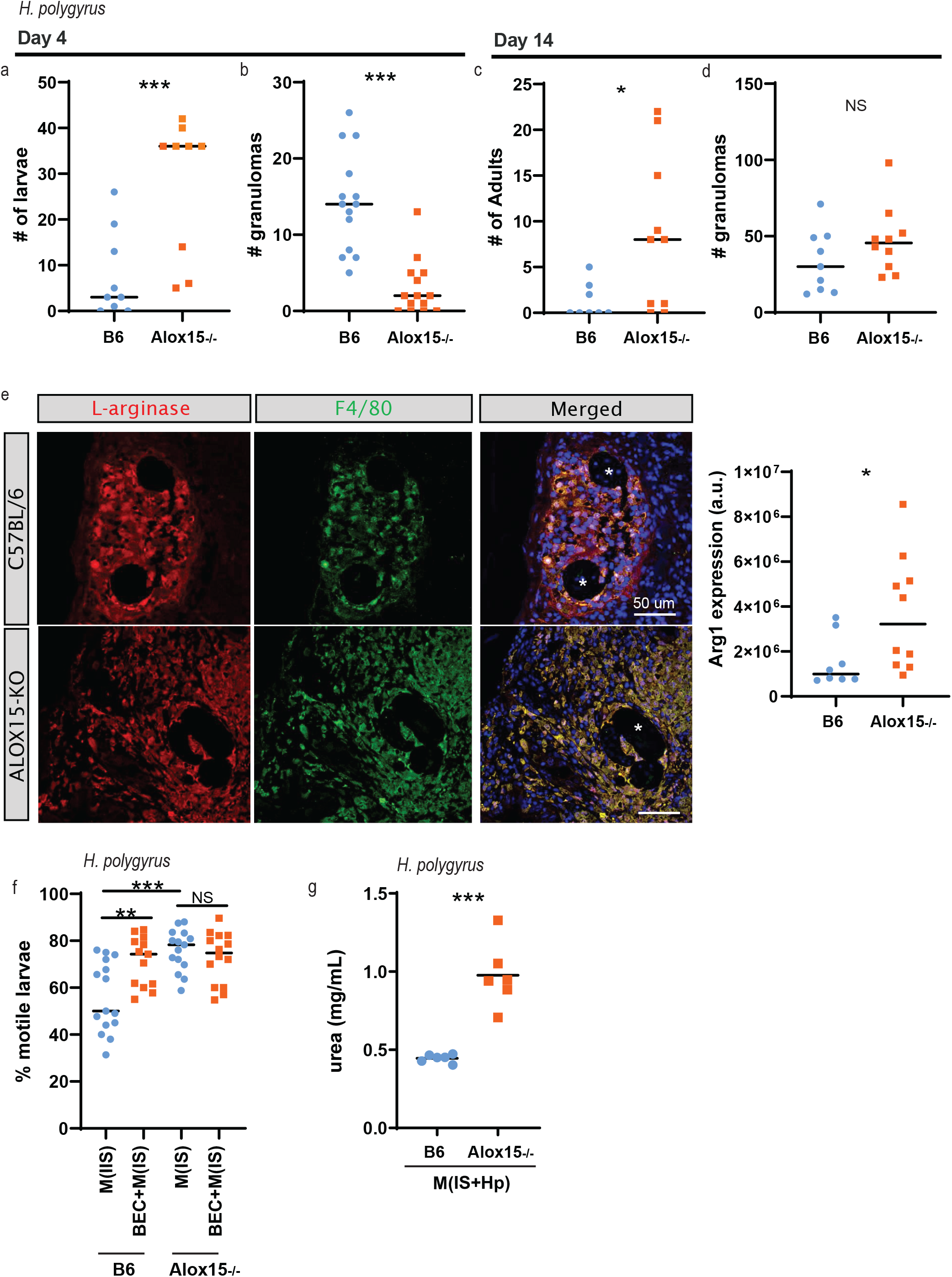
Alox15 is required for macrophage mediated protection against *H. polygyrus*. **A-D**. Alox15-/- and B6 controls were infected twice with 200 Hpb L3. 4 or 14 days after secondary infection, parasites and granulomas were enumerated from the small intestine. N=4-5 mice per group, 2 independent experiments. **E.** Immunofluorescence staining of Arginase and F4/80 in intestinal granulomas surrounding Hpb larvae at 4 days post-reinfection. N= 5-6, Quantification from 8-10 granulomas for each genotypes. **F.** Murine bone marrow-derived macrophages were grown from C57BL/6/J (B6) or ALOX-15^-/-^ mice. Differentiated BMDM were co-cultured with 100 third-stage Hpb larvae in presence of immune serum (IS) and with or without the arginase inhibitor BEC. The percentage of motile larvae was assessed for each genotypes by microscopy. n=3-9 mice, pool of 3 independent experiments. **G**. Arginase-1 activity of BMDM stimulated for 24 hours with IS and 100 Hpb L3 measured by urea production. (n=6, representative of 2 independent experiments).

To further generalize our findings, we took advantage of a recently published single cell RNA sequencing dataset comparing BALB/c versus B6 pleural immune cells in their response against the filarial pathogen *Litomosoides sigmodontis* [39]. The authors did show that in BALB/c susceptible mice, the presence of Type 1 cytokines prevent incoming monocytes to fully differentiate in tissue resident cells and to M2 shift. Interestingly, we found that Alox15 expression was enriched in tissue macrophages in resistant C57BL/6 mice after infection, but not in transitional hybrid M1/M2 macrophages in susceptible BALB/c mice (Figure S4b). Furthermore, like in the case of infection with Nb and Hpb, Arg1 and Rtnla expression were similar between the two macrophage populations.

Altogether, these observations show that Alox15 is a crucial enzyme involved in macrophage M2 commitment, thus affecting anti-helminth immunity.

### PPARδ activation can overcome Alox15 deficiency in helminth mediated trapping/killing

Alox15 is a lipid metabolizing enzyme, with a broad role in the production and metabolism of lipid mediators important in health and diseases [19]. Recently, Alox15 has been proposed to be a key metabolizing enzyme in T regulatory cell differentiation and regulatory phenotypes [40]. In absence of Alox15, Tregs had low FOXP3 expression and a high glycolytic profile. Supplementation with resolvins, lipid mediators downstream of DPA, EPA or DHA, rescued the altered T regulatory phenotype. Due to the parallels between our findings, we reasoned that Alox15 deficiency in M(IL-4) could be rescued by lipid supplementation.

To identify such lipids of interest, we performed lipidomic analysis of Hpb-infected intestines 4 days after secondary infection in WT and in Alox15-/-mice. Larval products of Hpb have previously been shown to trigger a broad anti-inflammatory eicosanoid shift after primary infection by suppressing the 5-lipoxygenase pathway, and inducing the cyclooxygenase (COX) pathway [41]. In secondary infection, arachidonic acid as well as linoleic acid were found in a similar quantity in WT and Alox15-/-animals (figure 8a). 12 and 15-HETE, two downstream products of Arachidonic acid were significantly decreased in Alox15-/-intestine as compared to WT counterparts. Similarly 17-HDHA, downstream of DHA was also reduced in Alox15-/-tissue. Although not significant, a similar trend could be observed for 9 and 13-HODEs downstream of Linoleic acid. We found no significant difference between genotypes in any other eicosanoids tested or in RvD1 (figure 8a).

**Figure 8:**
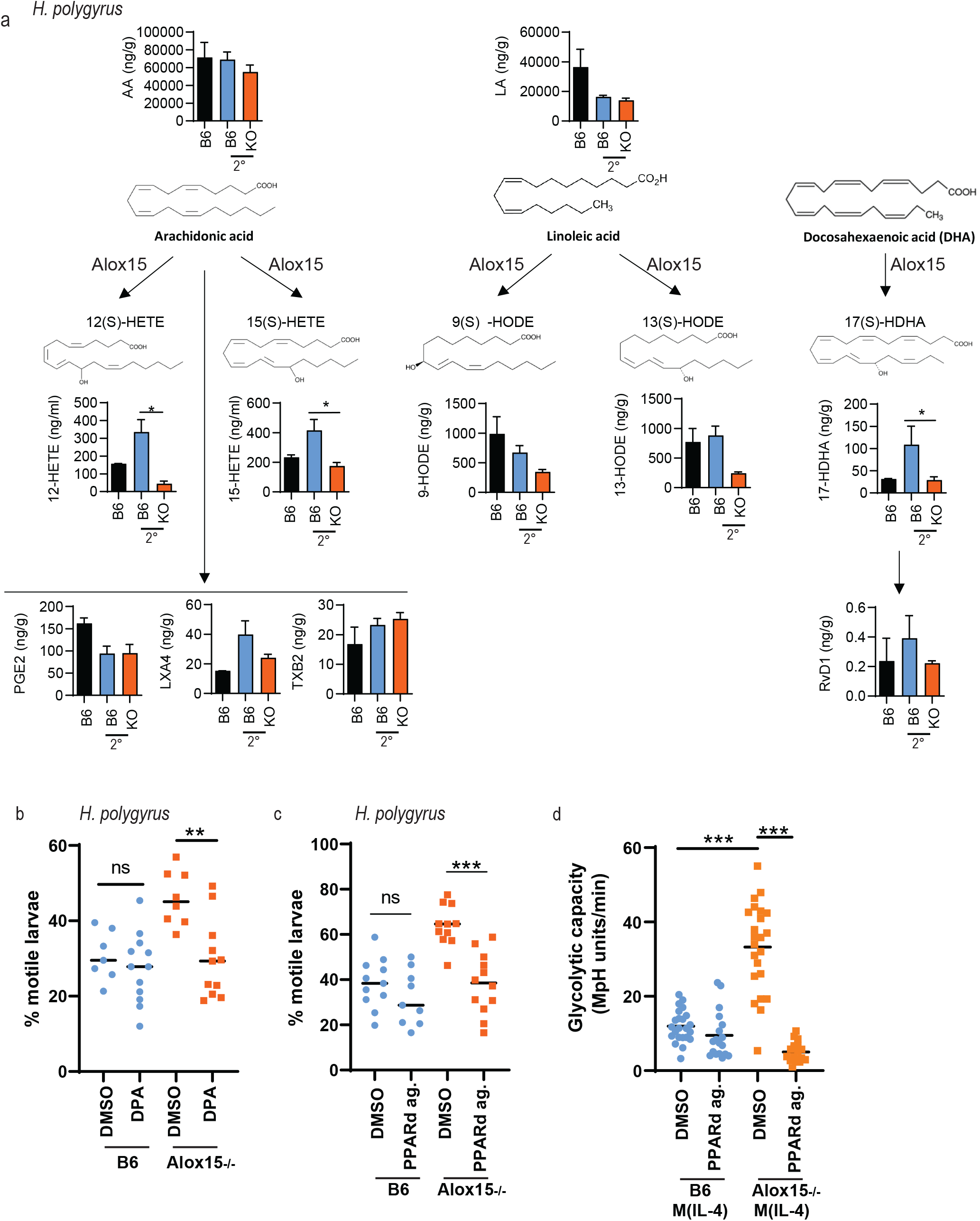
Heightened glycolysis cause loss of anti-helminth killing in Alox15-/-M2. **A.** LC-MS/MS analysis of LOX metabolites of Arachidonic acid (AA), of Linoleic acid (LA) and of Docosahexaenoic acid (DHA) in intestinal culture supernatants from Hpb infected mice N=4 per genotype, baseline for B6 N=2. **B&C.** Murine bone marrow-derived macrophages were grown from C57BL/6/J (B6) or ALOX-15^-/-^ mice. Differentiated BMDM were co-cultured with 100 third-stage Hpb larvae in presence of immune serum (IS) and with or without the Docosapentaenoic acid (DPA) or a PPARd agonist. The percentage of motile larvae was assessed for each genotypes by microscopy. N= 3-6 per condition, pool from 3 independent experiments. Of note B&C share some of the DMSO conditions. **D.** Glycolytic capacity of BMDM stimulated as with IL-4 and/or PPARd agonist for 24 hours prior to the assay. Data representative of 3 independent experiments, n = 5 wells per conditions, each conditions with 1-3 mice per experiments.

Omega-6 PUFAs are the most abundant polyenoic fatty acids in mammalian cells and are thus major lipoxygenase (LOX) substrates. We thus focused first on arachidonic acid and linoleic acid metabolism. Using the Hpb trapping assay, we performed rescue experiments by supplementing with 9-HODE, 13-HODE and 15-HETE the most common downstream mediator products of Alox15. However, none of those reversed the increase Hpb motility observed in co-culture assay with Alox15 deficient macrophages (Figure S5a).

Given we had also observed a decrease in Omega-3 (n-3) PUFA downstream products of Alox15, we further considered if those could be involved in our helminth-specific phenotype. It has recently been shown that n-3 DPA supplementation can enhance macrophage efferocytosis and phagocytosis [42]. We thus assessed if n-3 DPA supplementation could rescue the absence of trapping/killing observed in Alox15 deficient macrophages. Indeed, n-3 DPA supplementation rescued the trapping of Hpb larvae by Alox15-/-macrophages (Figure 8b). To confirm, this phenotype reversal was not specific to Hpb trapping or BMDM, we performed an *ex vivo* binding assay with IL-4 polarized lung macrophages and Nb larvae. Similarly, DPA was able to rescue the binding of Nb larvae by Alox-15 deficient macrophages (Fig S4B).

Given Omega-3 PUFAs and downstream metabolites are ligands, or partial-ligands, of peroxisome proliferator-activated receptors (PPAR), we next used an agonists of PPAR-γ and PPAR-δ, Rosiglitazone and GW501516 respectively as another method to validate that enhancing lipid metabolism could reverse the Alox15-/-phenotype. Interestingly, both PPAR agonists rescued the macrophage larvicidal activity against Hpb larvae (Fig S4C). GW501516, the PPAR-δ agonist led to a significant reversal in the impaired ability of Alox15-/-macrophages to reduce larval motility, similar to the trend observed with exogenous DPA (Figure 8c).

To further confirm that Alox15 downstream lipid metabolites control anti-helminth activity through PPAR mediated control of glycolysis, we performed Seahorse XF analysis of glycolysis and OXPHOS after rescuing Alox15-/-M(IL-4) with PPAR-δ stimulation. As expected both glycolysis (Figure 8d) and OXPHOS (Fig S4D) normalized to the level obtained for WT. Altogether our results show that Alox15 metabolism of DPA activates PPAR-δ that in turn controls glycolysis activation to ensure full M2 commitment.

## Discussion

Macrophages are central to anti-helminth immunity in the vast majority of mouse models but are largely unstudied in human helminthiasis. In rodents, macrophages require a so-called ‘alternative’ or “M2” activation mediated by type 2 cytokines or antibodies [4–6, 29, 34]. This M2 signature is also enhanced by collectins and stress signals [25, 30]. However, macrophage M2 polarization differs between mouse and human, at least in terms of hallmark genes. With the long-term goal of elucidating anti-helminth role of macrophage in humans, we investigated Alox15, an arachidonate lipoxygenase that is a hallmark of human macrophage alternative activation [12, 19]. Upon genetic deletion of Alox15, we found that macrophages cannot efficiently trap and kill hookworm larvae in mice, nor could they repair tissue damage. Interestingly, hallmark genes of alternatively activated macrophages were not compromised, begging the question as to why there was a loss of function. We further found that Alox15-/-macrophages activated with IL-4 displayed a hybrid phenotype of classical and alternative activation. This was associated with an uncontrolled switch to glycolysis. We finally show that to ensure a ‘pure’ M2 commitment, Alox15 derived metabolites of DPA are required to activate PPAR-δ, which keeps glycolysis in check.

For several decades, hallmark M2 genes have been used as a readout of alternative activation. Our study demonstrates that upregulation of such genes was not sufficient for functional activation; *i.e.* trapping helminth parasites or repairing tissue damaged by parasite’s migration. Notably, Arginase-1 has previously been proposed to participate in macrophage-mediated protection against helminths as both a marker and enzymatic effector molecule [4–6, 29]. Our observation that Arginase 1 activity did not functionally correlate with anti-helminth activity is in line with recent findings showing that macrophage-specific STAT6 activation is dispensable for trapping *H. polygyrus* in an arginase-independent manner [9]. Instead, Westermann and colleagues propose that intestinal epithelial cells make an important contribution to the formation of intestinal granulomas in a STAT6 dependent manner. However, the authors show that macrophage STAT6 expression is still required for protection against *N. brasiliensis* in line with our previous findings [4]. Here, we found that Alox15 expression in macrophages was required for both *H. polygyrus* and Nb trapping *in vitro.* We further confirmed *in vivo* that macrophage deficiency solely in macrophage is sufficient to cause a loss of protection.

For both Nb and Hpb, macrophages binding to the larvae was compromised by Alox15 deficiency, suggesting that parasite recognition might be affected by Alox15 deficiency. However, comparing Arginase-1 blockade (by BEC) with Alox15 genetic deficiency did not lead to any common genes downregulated. It is important to note here that this transcriptomic analysis was performed after 24 hours of IL-4 stimulation, but Alox15 expression is known to peak at 48 hours, therefore it is possible we simply missed the optimal window to assess this response [43].

*N. brasiliensis* infection in mice typically gives a rapid, ‘pure’ Th2 response. This textbook anti-helminth immunity is distinct from that observed outside the laboratory, as human or wild animal helminthiasis is characterized by a ‘modified type 2’ response thought to be a hybrid of Th2/Th1/ Tregs [44, 45]. Although there are discrepancies in the literature, chronic human hookworm infection has been associated with a mixed cytokine response; IFNg, TNFa, and IL-10 are elevated in the serum in addition to IL-5 [46–48]. Recently, CyToF of helminth-exposed patients has deepened our understanding of the immune response in humans and confirmed this hybrid response is due to ILC2s, and CD4+, CD8+, and γδ T cells [46]. To date, we could not find any reports of mixed polarization of macrophages in helminth infected individuals, however this is likely due to the fact that peripheral responses are studied in the context of human helminth infection. In future, flow cytometry of, for example, sputum samples from infected patients could give insights into macrophage polarization. We speculate that in this mixed cytokine environment, macrophages might not commit to a purely ‘alternative’ phenotype, and instead develop a hybrid M1/M2 commitment as recently shown for sequential IL-4/LPS stimulation [49]. Given the central role of M2 macrophages in protection against reinfection in rodent helminth models, such a defect in M2 polarization could explain why hookworm-infected individuals are highly susceptible to re-infection for life. In line with this idea, Finlay et al. recently compared two mouse genotypes with different susceptibility to filarial infection and reported that in the presence of Type 1 cytokines, tissue resident macrophages do not properly adopt the M2 phenotype, and instead display lower OXPHOS and heightened phagocytosis. [39] Interestingly, despite an overall decrease in an IL-4-assosciated gene signature, the authors report that the hallmark M2 genes were not strongly affected, similar to our model. By reanalyzing the single cell sequencing data published by Finlay and colleagues, we have further shown that Alox15 expression, but not Arginase 1, was correlated with parasite clearance and efficient M2 polarization, highlighting that Alox15 is likely to be a generic anti-helminth feature of macrophage polarization.

As highly plastic cells, the polarisation of macrophage activation has been described as a continuum with ‘classical’ and ‘alternative’ states at either end. While comparisons of *in vivo* and *in vitro* datasets have begun to challenge the simplicity of this model [33, 50], the terms are still widely used. It has been proposed in recent years that classically activated macrophages enhance their glycolysis to meet their energetic need, while alternatively activated macrophages use the more energy efficient oxidative phosphorylation. Yet there is some conflict over the requirement for glycolysis in alternative activation [33, 50], which is thought to be due, in part, to the use of the inhibitor 2-DG to block glycolysis; 2-DG appears to have off target effects on OXPHOS activity. There is reason to suspect that intact glycolysis is required for efficient anti-helminth immunity. For example, inhibiting glucose utilization by intraperitoneal injection of 2-DG during *H. polygyrus* infection perturbs M2 polarisation, significantly reducing expression of RELMa [51]. Furthermore, it has been noted that alveolar macrophages are not protective against the lung phase of *N. brasiliensis* infection as they are hyporesponsive to IL-4 due to the glucose-restricted environment of the alveolar niche [32].

While M1 polarization clearly requires metabolic reprogramming, M2 polarization may be more subtle. In our own study, we noted that OXPHOS activity was not different between basal ‘M0’ and IL-4 activated ‘M2’ bone marrow-derived macrophages. Here, we argue that glycolysis needs to be kept in check to obtain a functional M2 commitment. Failing in that control results in the development of a hybrid M1/M2 signature which lacks anti-helminth activity, but retains M1 functions, such as anti-*Leishmania* control. This suggests that alternative effector functions are more tightly regulated than classical effector functions. In line with our work, a hybrid M1/M2 signature has recently been reported after sequential exposure of BMDM to IL-4 and LPS *in vitro*, with a metabolic profile matching the one we described for Alox15-/-M(IL-4), *i.e.* a heightened glycolysis at the same time as a heightened OXPHOS[49]. The authors then prove that the pro-inflammatory genes expressed by hybrid M1/M2 macrophages require HIF-1α stabilization as in the case of ‘classical’ polarization. Whether this also occurs in the absence of Alox15-/-should be further investigated. Other hybrid M1/M2 macrophage phenotypes have recently been reported in response to IL-33 stimulation which elicits the co-expression at the protein and mRNA level of alternative markers (Arg1, Retnla, Chil3, Alox15…) as well as pro-inflammatory markers (IL-6, IL-1b) [52]. Interestingly, IL-33-induced hybrid polarization did not display enhanced glycolysis but rather full integrity of the TCA cycle. IL-33 is very strongly released in the lungs after Nb infection and given we observed stronger transcriptional differences *in vivo* that *in vitro* between WT and Alox15-/-macrophages, it could be the case that Alox15 is also important to the fine tuning of IL-33 activation of macrophages.

Despite a staggering number of publications on Alox15, its role in cellular metabolism has not been well-studied to date. Namgaladze et al. first established the possible involvement of Alox15 in cellular immunometabolism in 2015 [53]. Indeed, the authors showed that AMPK inhibition blocked Alox15 upregulation after IL-4 stimulation, but did not further characterize the activation profile of said macrophages. Instead, they investigated the role of AMPK and Alox15 in the IL-4 priming effect prior to LPS activation. In this context, they could show that Alox15 deficiency decreased the pro-inflammatory signature of IL-4 primed macrophages. Treatment with 12-or 15-HETE instead of IL-4 could similarly potentiate pro-inflammatory cytokine production in macrophages. In our model, 12- and 15-HETE supplementation was not sufficient to restore the anti-helminth activity of Alox15-/-M(IL-4). We have unfortunately not assessed cytokines production after this supplementation. However, n-3 DPA supplementation, another lipid substrate for Alox15, was able to restore a ‘pure’ M2 commitment in Alox15-/- M(IL-4), both functionally and metabolically. A current limitation of our study is that we have not identified the responsible downstream metabolite(s) of DPA.

Interestingly a similar finding has been reported in regulatory T cells[40], where Alox15 deficiency has been shown to dampen Treg activation and cause upregulation of pro-inflammatory factors. Supplementation with resolvins –downstream products of Alox15 activity on DPA/DHA – was shown to rescue Treg activation. Of note, the functional relevance of resolvins is a highly debated topic in the field due to their production in very low quantity and the associated challenge of detecting them with current technology. Using metabolomics, the authors reported heightened glycolysis and TCA in Alox15-/-Tregs. Together with our results, this suggests that the SPMs/Alox15 pathway might be crucial as a ‘confirmation signal’ to allow the development of a pro-repair, anti-helminth M2 commitment. Failure to deliver this secondary signal may then push the immune cell towards a pro-inflammatory phenotype as a ‘fail-safe’ or ‘default’ pathway.

## Methods

### Reagents

The following reagents were used in various assays: IL-4 10 ng/mL (Peprotech), LPS 10 ng/ml (Peprotech), IFNg: 10 ng/mL (Peprotech) and were reconstituted in PBS.

BEC 1 uM (Sigma), Rosiglitazone 0.1, 0.5, 1uM (PPAR-γ agonist, Sigma), GW501516 0.1, 1, 5uM (PPAR-δ agonist, Sigma), n-3 Docosapetaenoic acid sodium salt DPA 100uM (Cayman Chemical) 15-HETE (Cayman Chemical), 13-HODE (Cayman Chemical), 9-HODE (Cayman Chemical) were reconstituted in DMSO (Sigma).

### Mice

C57BL/6J (WT) mice were obtained from Charles River Laboratories (Germany) or from Animal Resources Center (Australia). All mice were maintained under specific pathogen-free (SPF) conditions at École Polytechnique Fédérale de Lausanne (EPFL), Switzerland or by the Monash Intensive Care Unit Facility at Monash University, AMREP campus Melbourne, Australia or by the Animal Facility, Swiss TPH, Switzerland. All mice were age and sex-matched and used between 8-14 weeks of age. Littermates of the same sex were randomly assigned to experimental groups. Mice were maintained at 3-5 animal per cage and ad libidum access to water and food. All animal experiments were approved by the Service de la consommation et des affaires vétérinaires (1066 Épalinges, Canton of Vaud, Switzerland) with the authorization number VD 3001, or by the AEC committee of the Alfred campus, Melbourne Australia with authorization number E/1846/2018/M or by BL/543/2022.

### Preparation and Isolation of *N. brasiliensis* Larvae

Nb was maintained by monthly passage in Lewis rats. Infective larvae (L3) were prepared from 2 week old rat fecal cultures as previously described (Camberis et al., 2003). 550L3 or 250 L3 were delivered by subcutaneous injection in 100 μL of sterile PBS to the scruff of the neck.

Viable larvae were recovered from tissue (lung or gut) by baerman at 37 °C for at least 2 h as previously reported (my method paper). Viable worms follow a thermal gradient and migrate out of the bag to collect at the bottom for counting. Larvae present in the large airways were harvested by bronchoalveolar lavage, with 3 × 1 ml of PBS, followed by filtering on 40 um gause. For larvae count and red blood cell count, the 3ml of wash were pooled together. Larvae were manually counted under brightfield on a stereomicroscope. Red blood cells were counted with a haemocytometer, after centrifugation of the BALF and resuspended in 1 ml to account for the difference of BALF volume harvested in each mouse. D0 marks the day of infection. If not explicitly mentioned, the time between primary and reinfection was 30 days.

For adoptive transfer, 1.7-2 million cells (stimulated for 24 hrs with IL-4) were administered intranasally in 50 μl of PBS, the evening prior to infection with 550L3.

### Preparation and Isolation of *H. polygyrus bakeri* Larvae

The standard Hpb infective stage-three larvae (L3) were generated in the laboratory based on published protocols [54]. 200 L3 were administered per os as previously described [6]. Viable worm burden were obtained by baerman as previously described [55]. Granulomas were enumerated by manual counting of the small intestinal contents under a dissecting microscope. D0 marks the day of infection. If not explicitly mentioned, the time between primary and reinfection was 30 days.

### *Leishmania mexicana* maintenance and infection

Transgenic *L. mexicana* stably expressing the fluorescent protein TurboRFP (L. mexicanaTurboRFP) were maintained in promastigote stage at 27°C in RPMI media pH 7.4 supplemented with 10% FCS with biweekly passaging. Only cultures <20 passages since isolation from infected murine tissue were used to generate amastigotes for infections. Amastigotes were generated by incubating stationary phase promastigotes in RPMI media pH 5.5 supplemented with 20% FCS for 4 days at 33°C 5% CO2.

### Macrophage generation

For bone marrow-derived macrophages, marrow was flushed from the femur and tibia of C57BL/6 mice and passed through cell strainers (70 µm) in PBS. The cells (10^6^ mL^−1^) were cultured in 30% M-CSF (supernatant of L929 cell culture) supplemented medium (RPMI (Gibco), 10% FCS (Gibco), penicillin/streptomycin) for 7 days as described previously [34]. Macrophages were harvested between day 7 and 10 of culture for experiments.

For lung macrophages, mice were either used naïve or infected with 250 or 550 L3 subcutaneously. The lungs were washed by bronchoalveolar lavage with 3 mL of PBS and then digested in media (DMEM (Gibco), 2 mg mL^−1^ collagenase (Sigma), DNase 12.5 U mL^−1^ (Roche, Sigma-Aldrich, Merck & Cie, Schaffhausen, Switzerland)) at 37°C on an orbital shaker for 45 min. The cells were passed through a cell strainer (70 µm) and seeded (2.5 × 10^5^) directly in a flat-bottom 96-well plate. Non-adherent cells were washed away the following day and adherent macrophages were cultured in complete DMEM (10% FCS, 1% penicillin/streptomycin) for 2 days before co-culture with larvae.

### Larval binding assay

<u>Nb assay</u>: macrophages were polarized or not with recombinant IL-4 (10 ng mL^−1^) and various inhibitors or agonists (see reagents) for 24 hours. 0.1×10^6^ cells were then co-cultured with 100 third-stage *N. brasiliensis* larvae (Nb L3) in 96-well plates, and the number of larvae with >10 cells attached to their surface was manually counted by light microscopy after 4 days of co-culture [56], and expressed as a percentage of total larvae per well.

<u>Hpb assay:</u> 0.1×10^6^ BMDM were co-cultured with 100 third-stage *H. polygyrus bakeri* (Hpb) in presence of immune serum. Inhibitors, supplements or agonists were provided 24 hours prior to adding the larvae. All co-cultures were performed at 37°C, 5% CO_2_, for 24 h. The percentage of motile larvae attacked by macrophages were then quantified using a stereo microscope.

### Larval viability assessment

After the co-culture, Sytox green was added (1:100, Life technologies, Cat#S7020) to the wells and viability of the larvae was assessed using fluorescent microscopy. Larvae were considered dead, or to have impaired viability, when staining positive for the dye, and expressed as a percentage of total larvae per well.

### Leishmania/macrophage co-culture assays

BMDM infection with *L. mexicanaTurboRFP* and subsequent flow cytometric analysis were performed as described previously [57]. Briefly, BMDMs were seeded into non-treated 96-well plates at 110,000 cells/well. Following overnight incubation at 33°C, BMDMs were co-incubated with *L. mexicanaTurboRFP* amastigotes at a low (0.5:1) or high (3:1) multiplicity of infection at 33°C 5% CO2 for the indicated period. At the indicated time point, infected macrophages were gently detached from plates and single cell suspensions were stained with a fixable viability dye, anti-F4/80 and anti-CD11b, and analyzed by flow cytometry. Dead cells and extracellular parasites were excluded prior to gating of infected macrophages (SSChiFSChiF4/80+CD11b+TurboRFP-H+) and uninfected macrophages (SSChiFSChiF4/80+CD11b+TurboRFP-H-). Data was analyzed using FlowJo software where infected macrophages are expressed as the percentage of gated viable, single macrophages. The TurboRFP geometric mean fluorescence intensity (gMFI) of infected macrophages minus that of uninfected macrophages was used as an indicator of average parasitic burden. Parasitic index, the product of the percent infection and parasitic burden, was used as a more general readout of infection.

### RNA isolation and RT-qPCR

RNA of 1 × 106 cells was extracted with a TriPure (Sigma-Aldrich), and reverse transcribed using FIREScript RT cDNA synthesis Kit Solis Biodyne (Lucerna 06 −15-00200) for qPCR analysis. qPCR was performed using PowerUp SYBR green Applied Biosystems (Thermofisher A25741) on an Bio-Rad CFX96 touch Real-Time PCR detection system. Data analysis was performed with CFX Maestro, and expression was normalized according to expression of the housekeeping gene GAPDH.

### Multiplex PCR

Custom oligonucleotide primers and probes (Table 5) were designed using National Center for Biotechnology Information (Bethesda, Md) primer blast software and were purchased from Microsynth (Balgach, Switzerland). Amplification was carried out using iQ Multiplex Powermix and a CFX96 Real-Time detection system using forward and reverse primers at a concentration of 10μM and probes at a concentration of 50μM. Absolute quantification was performed using the CFX Manager software 2.1 (all from Bio-Rad) based on values obtained with a set of purified amplicons used as standards.

**Table 5.**
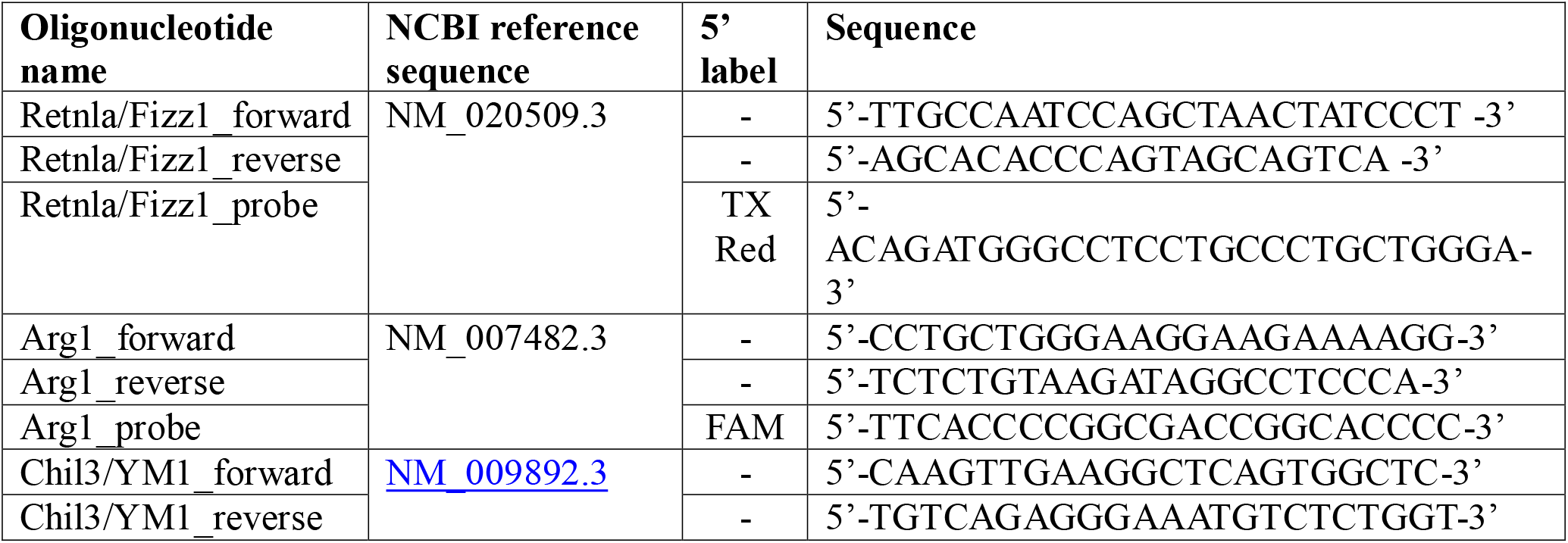
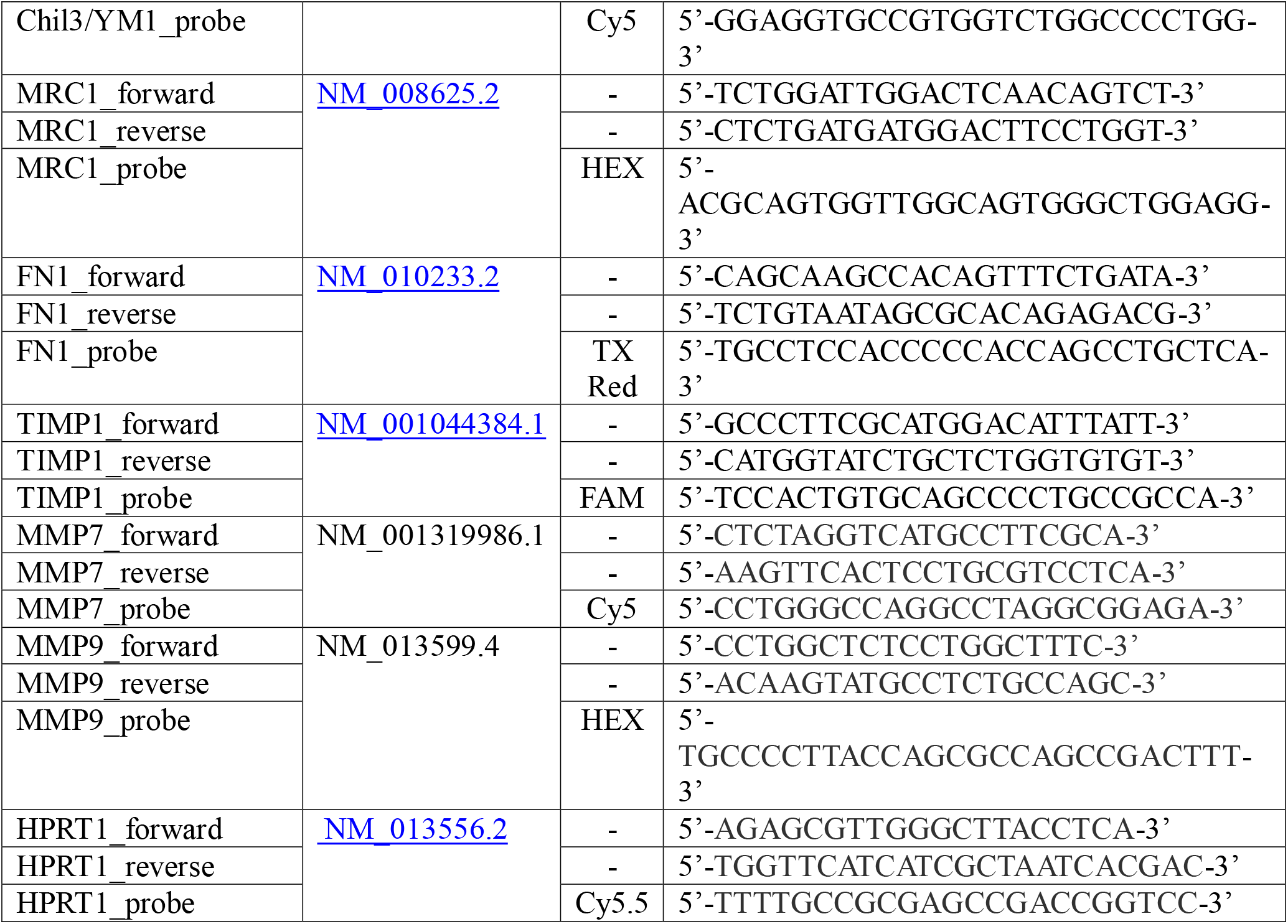
Oligonucleotide primers and probes for multiplex PCR analysis.

The primers and probes specific for Retnla/FIZZ1, ARG1, CHIL3/YM1, MRC1; and FN1, TIMP1, MMP7, and MMP9; were collectively used in multiplex-1, and −2 respectively. The primers and probe specific for HPRT1, our reference gene, were added into each multiplex. *Cy*, Cyanine; *FAM*, <u>fluorescein</u>; *HEX*, hexachloro-fluorescein; *Tx Red*, Texas Red.

### BMDM transcriptomics

Fastq files were processed using the Laxy tool [58] using the RNAsik pipeline [59] using STAR aligner [60] with Mus musculus GRCm38. Reads were quantified using featureCounts [61] producing the raw genes count matrix and various quality control metrics, summarised in MultiQC report [62]. The gene count matrix was analysed with Degust [63], a web tool which performs differential expression analysis using limma voom normalisation producing counts per million (CPM) library size normalisation and trimmed mean of M values (TMM) normalisation for RNA composition, and also several quality plots such as classical multidimensional scaling [MDS] and MA plots. Samples were batched corrected to account for known variation across replicates. This was done within Degust by adding a covariate to the linear model for each sequencing batch which reduced the noise and improved the power of the test. Differentially expressed genes were defined as those showing a >2-fold (log2FC 1) change in expression with a false-discovery rate (FDR) of 0.05.

### Single cell sequencing

Lungs were minced and then digested in media (DMEM (Gibco), 2 mg mL−1 collagenase (Sigma), DNase 12.5 U mL−1 (Roche, SigmaAldrich, Switzerland)) at 37°C on an orbital shaker for 45 min at 105 rpm. Lungs were then passed through a 70µm cell strainer with syringe plungers. Red blood cell lysis was performed (Merck, 11814389001). Whole transcriptome analyses of FACS sorted CD45+ cells were performed using the BD Rhapsody Single Cell Analysis System (BD, Biosciences). Cells were stained with Zombie Aqua live/dead (Biolegend) and Fc block (2.4G2, BioXCell, BE0307) according to maufacturer’s instructions. Prior to FACS staining, cells were labelled with sample tags (633793 BD Mouse Single-Cell Multiplexing Kit) according to the manufacturer’s protocol. As the CD45 sample tag clone and FACS antibody clone were the same (30-F11), sample tagging was performed in the presence of a 1:1 (w/w) quanitity of anti-CD45 FACS antibody. In brief, for each condition, 10^6^ cells were resuspended in staining buffer (1% BSA, 1% EDTA in PBS) and incubated with the respective Sample Tag:anti-CD45 BUV395 cocktails for 20 min on ice. Cells were then transferred to a 5-ml polystyrene tube, washed twice with 2 ml staining buffer and centrifuged at 400g for 5 min. Samples were stained in a FACS antibody cocktail mix for 20 mins on ice: CD19-APCe780 (1D3, ebioscience, 47-0193-80), Gr-1-PE (RB6-8C5, Biolegend, 108408), SiglecF-PE-CF594 (E50-2440, BD, 562757), Ter-119-Biotin-SA-PECy5 (Biolegend, 116204), CD11c-Pecy7 (HL3, BD, 564079), CD3e-BUV737 (145-2C11, BD, 564618), CD11b-PacBlue (M1/70, Biolegend, 101224), CD4-BV711 (RM4-5, eBioscience, 14-0042-83),CD8-BV786(H35-17.2, BD, 740952). Cells were washed and resuspended in 1 ml staining buffer for counting. Next, an equal number of cells from 4 barcoded samples per group were pooled and the mixture was centrifuged at 400g for 5 min. The BD Rhapsody cartridge was loaded with 40,000 cells. Single cells were isolated with the BD Rhapsody Express Single-Cell Analysis System according to the manufacturer’s recommendations (BD Biosciences). cDNA libraries were prepared using the BD Rhapsody Whole Transcriptome Analysis Amplification Kit (633801 BD Biosciences) following the BD Rhapsody System mRNA Whole Transcriptome Analysis (WTA) and Sample Tag Library Preparation Protocol (BD Biosciences). The final libraries were quantified using a Qubit Fluorometer with the Qubit dsDNA HS Kit (Q32851 Thermo Fisher Scientific). Library size distribution was measured with the Agilent high-sensitivity D5000 assay on a TapeStation 4200 system (5067-5592 Agilent Technologies). Sequencing was performed in paired-end mode (2 × 75 cycles) on a NovaSeq 6000 with NovaSeq 6000 SP Reagent Kit chemistry.

### Single cell sequencing analysis

Gene quantification was generated with STARsolo (STAR-2.7.10a, [64]) and aligned to a GENCODE Version M29 reference [65] with sample tags added for de-multiplexing purposes. Parameters were in accordance with DB Rhapsody (version 1) with default settings. Subsequently, the resulting filtered matrix underwent quality control assessments and analytical evaluations within the R environment (version 4.3.2, [66]), utilizing various R packages such as SoupX (version 1.6.2, [67]) for the removal of ambient mRNAs, scater (version 1.30.1, [68]) to identify outlier metrics and Seurat (version 5.0.1, [69]). No other cell filtering criteria were applied regarding total number of UMIs, unique expression genes, or mitochondrial gene proportion. However, genes with cell counts less than 3 were excluded. Seurat’s demultiplexing function was employed to eliminate doublets and ambiguous cells. Additionally, Seurat’s SCTransform was applied using the ‘glmGamPoi’ method, followed by RunPCA, RunUMAP, FindNeighbors and FindClusters for clustering analysis. Default parameters were utilized, except for employing 40 dimensions of input features and proceeding with a resolution of 2 for cluster analysis. Manual annotations were conducted by referencing markers from various published sources [70–76]. UMAP figures were customized using ggplot2 (version 3.4.4, 14) [77].

### Sample preparation for LC-MS/MS

For intestinal tissue culture, the intestine was removed, extensively flushed with cold PBS. The intestine was isolated, cut longitudinally and placed separately in 100 mm Petri dishes with PBS containing 10 µg/ml Gentamicin, 100 U/ml Penicillin and 100 µg/ml Streptomycin (all from GIBCO). Opened cut intestines were gently scraped to remove mucus, washed 3 times and transferred into a new Petri dish. The anterior 4 cm of the intestine (duodenum) were cut into 1–2 mm pieces and transferred to a 24-well plate with 2 ml/well of RPMI media (PAA Laboratories) containing 10% heat inactivated FCS, 10 µg/ml Gentamicin, 100 U/ml Penicillin and 100 µg/ml Streptomycin (all from GIBCO). 24-well plates were incubated over night at 37°C before supernatants were harvested and analyzed by LC-MS/MS.

Samples were prepared as triplicates in medium/MeOH (1:1) or PBS/MeOH (1:1) with an analyte concentration of 0.1, 1, or 10 ng/mL (10× higher concentrations for PUFAs). Automated solid phase extractions were performed with a Microlab STAR robot (Hamilton, Bonaduz, Switzerland). Prior to extraction, all samples were diluted with H_2_O to a MeOH content of 15% and 10 μL of IS stock solution was added. Samples were extracted using Strata-X 96-well plates (30 mg, Phenomenex, Aschaffenburg, Germany) and eluted with MeOH. Samples were evaporated to dryness under N_2_ stream and redissolved in 100 μL MeOH/H_2_O (1:1).

### LC-MS/MS lipid mediator analysis

Chromatographic separation of eicosanoids was achieved with a 1260 Series HPLC (Agilent, Waldbronn, Germany) using a Kinetex C18 reversed phase column (2.6 μm, 100 × 2.1 mm, Phenomenex) with a SecurityGuard Ultra Cartridge C18 (Phenomenex) precolumn. The Sciex QTRAP 5500 mass spectrometer (Sciex, Darmstadt, Germany), equipped with a Turbo-V^TM^ ion source, was operated in negative ionization mode. Identification of metabolites was achieved via retention time and scheduled multiple reaction monitoring (sMRM). Unique Q1/Q3 transitions were selected for each analyte by using single analyte injections and comparison with the literature. Analytes with identical MRM transitions were differentiated by retention time as described in [18]

### Flow cytometry

BMDM were polarized or not with recombinant IL-4 (10 ng mL^−1^, peprotech) for 24 hours. The next day, cells were detached with cold PBS and a cell scraper. Isolated cells were resuspended in FACS buffer, and Fc-receptors were blocked (2.4G2) before staining with monoclonal antibodies before collection on a Fortessa flow cytometer (BD). Fluorochrome-conjugated antibodies to the following cell surface molecules were used: CD45-AF700 (30-F11, 1:100, BD Biosciences Pharmingen), F4/80-APCeF780 (BM8, 1:100, Ebiosciences), Arginase-APC (1:100, Abcam), Ki-67-PE (SolA15, 1:10, Ebiosciences). Cell suspensions were stained with LIVE/DEAD fixable aqua dead cell Stain kit (Invitrogen) when fixation was required for intracellular staining. Analysis was performed with the FlowJo software (Treestar).

### Arginase-1 activity assay

Macrophage arginase-1 activity was determined according to previously published methods [6]. Briefly, 300000 adherent macrophages (48 well plate) were lysed and conversion of L-arginine was quantified indirectly by measuring urea production with a QuantiChrom Urea Assay (BioAssay Systems, Hayward, CA).

### Griess assay

Culture media from L. mexicanaTurboRFP infection assays was harvested by centrifuging to remove cells and retaining supernatant. 50 uL supernatant was first incubated at room temperature for 7 min with 50 uL 0.1% sulfanilamide in phosphoric acid prior to addition of 50 uL N-1-napthylethylenediamine dihydrochloride (NEDD) and a further 7 min incubation. Plates were then analyzed by plate reader at 540 nm. Sample nitrite concentration was determined by a nitrite standard dilution series.

### Fluorescence imaging

BMDM were grown on 8 chambers ibidi slides, fixed in 2% PFA. Prior to staining, cells were permeabilized in PBS, 0.5% triton X100 (Sigma) and blocked in 5% PBS, BSA (Sigma). stained with DAPI and anti-Alox15 antibody (1:100, Abb244205, Abcam).

Granulomas were identified on paraffin sections by light microscopy and serial sections were used for immunofluorescence staining for Arginase-1-AF568 (1:800) and F4/80-AF647 (BM8, 1:300, Ebiosciences) and DAPI (Sigma). Stained tissue sections were imaged with an inverted point scanning confocal microscope (Zeiss LSM 710) with a Plan-Apochromat (63×/1.4 NA or 40×/1.3 NA) objective.

Alox15 or Arginase intensity was measured in Fiji. To control for differences in background fluorescence between experiments and antibody/dye batches, the contrast was adjusted to minimize <u>autofluorescence</u> and a minimum brightness threshold was set such that only positive staining could be visualized. The same contrast and threshold values were applied to all images taken across all treatment groups within a single experiment using Fiji [78].

### Seahorse flux analysis

BMDM were polarized or not with recombinant IL-4 (10 ng mL^−1^, peprotech) for 24 hours and with LPS (10 ng/ml, Sigma) for 8 hours. The next day, cells were detached with cold PBS and a cell scraper. Isolated macrophages from individual mice or a pool of 2-3 mice were plated at 50 000 cells per well in 96 well plate (100 ul) or at 150 000 cells per well in a 24 well plate (450 ul) in XF media and allowed to adhere for 30 min at 37C, 5%C02. The cells were then transferred for 30 min to a 37C incubator without C02 just prior to analysis. ECAR and OCR were measured in XF media under basal conditions, in response to 25 mM glucose, 1.5 μM oligomycin, 1.5 μM FCCP, 1mM Sodium Pyruvate, 0.5 μM Antimycin A, 0.5 μM Rotenone (Sigma) using a 96-or 24-well extracellular flux analyzer XFe-96 and XFe-24 (Seahorse Bioscience) as previously described [79].

## Data analysis

All data were analyzed using GraphPad Prism 8.2 (GraphPad Software, La Jolla, CA, USA) or R 3.4.3.21. For LC-MS/MS analysis, all samples were normalized to their RNA content. Data were analyzed using Wilcoxon, Friedman, or Kruskal-Wallis test with respective post hoc test as appropriate. Significant differences were defined at P <0.05.

## Acknowledgements

We thank the MICU animal facility at Monash AMREP; the Monash Micro Imaging platform for support with imaging; Micromon for RNA sequencing. We thank the École Polytechnique Fédérale de Lausanne (EPFL) animal facility and sequencing facility. We thank Monash bioinformatic platform for performing analysis on the BMDM data set and would like to acknowledge that Nick Wong has done the bulk of the work on the BMDM dataset. This work was supported by the Swiss National Science Foundation (<u>SNF310030_156517</u>), and by a Monash Platform Access Grant (PAG19-0569). N.L.H. is supported by a National Health and Medical Research Council (NHMRC) of Australia SRF-B fellowship. T.B. is supported by a SNF-PRIMA Fellowship (<u>PR00P3_193084</u>). We thank Mark Haid and Jerzy Adamski for their support with metabolics studies.

## Author contributions

T.B., N.L.H. and J.E. conceived of the idea. T.B., N.L.H., J.E. and R.D. participated in the writing of the paper. R.D., T.B., M.M., C.G., V.B. conducted *in vivo* and *in vitro* experiments and downstream analysis. W.L. performed histology analysis. T.V. and U.N. performed qPCR analysis. H.F. performed lipidomics. C.R. performed sequencing of the single cell. B.A. conducted the analysis of the single cell dataset, both under M.B. supervision. S.A. performed the analysis of the lung RNA sequencing dataset. D.D. complemented the analysis of late Nick Wong at the Monash Bioinformatic platform for the BMDM RNA sequencing dataset. M.E.N.S and B.K performed the work on *Leishmania*. All authors discussed the results and commented on the paper.

## Materials & Correspondence

Correspondence and material requests should be addressed to.

