## supplemental figures for "Holding glycolysis in check though Alox15 activity is required for macrophage M2 commitment and function in tissue repair and anti-helminth immunity"

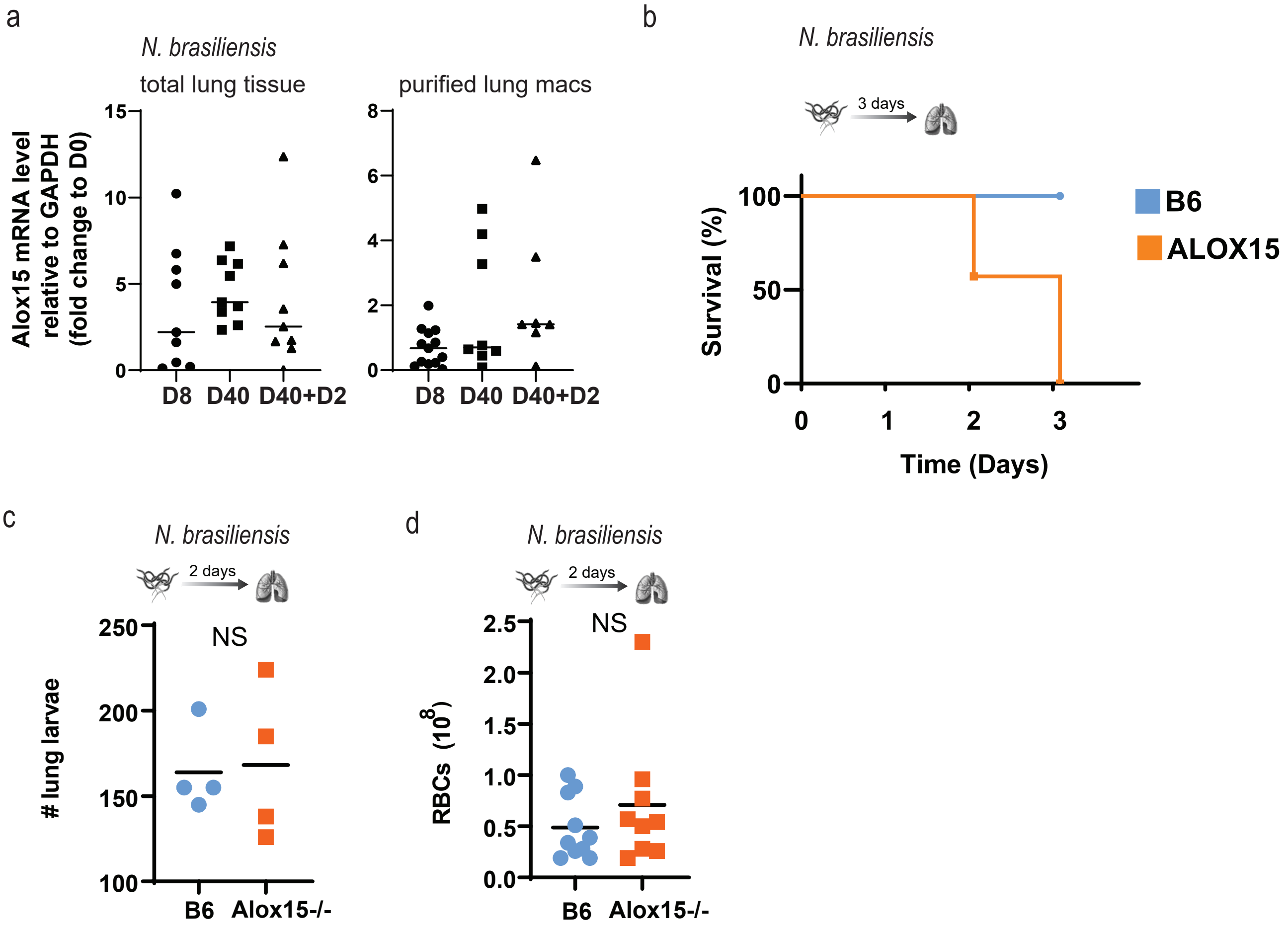

a
N. brasiliensis
total lung tissue
purified lung macs
Alox15 mRNA level relative to GAPDH (fold change to D0)
15
10
5
0
8
6
4
2
0
D8
D40
D40+D2
D8
D40
D40+D2
b
N. brasiliensis
3 days
Survival (%)
100
50
0
0
1
2
3
Time (Days)
B6
ALOX15
c
N. brasiliensis
2 days
NS
### lung larvae
250
200
150
100
B6
Alox15-/-
d
N. brasiliensis
2 days
NS
RBCs (10^8)
2.5
2.0
1.5
1.0
0.5
0.0
B6
Alox15-/-


Figure S1

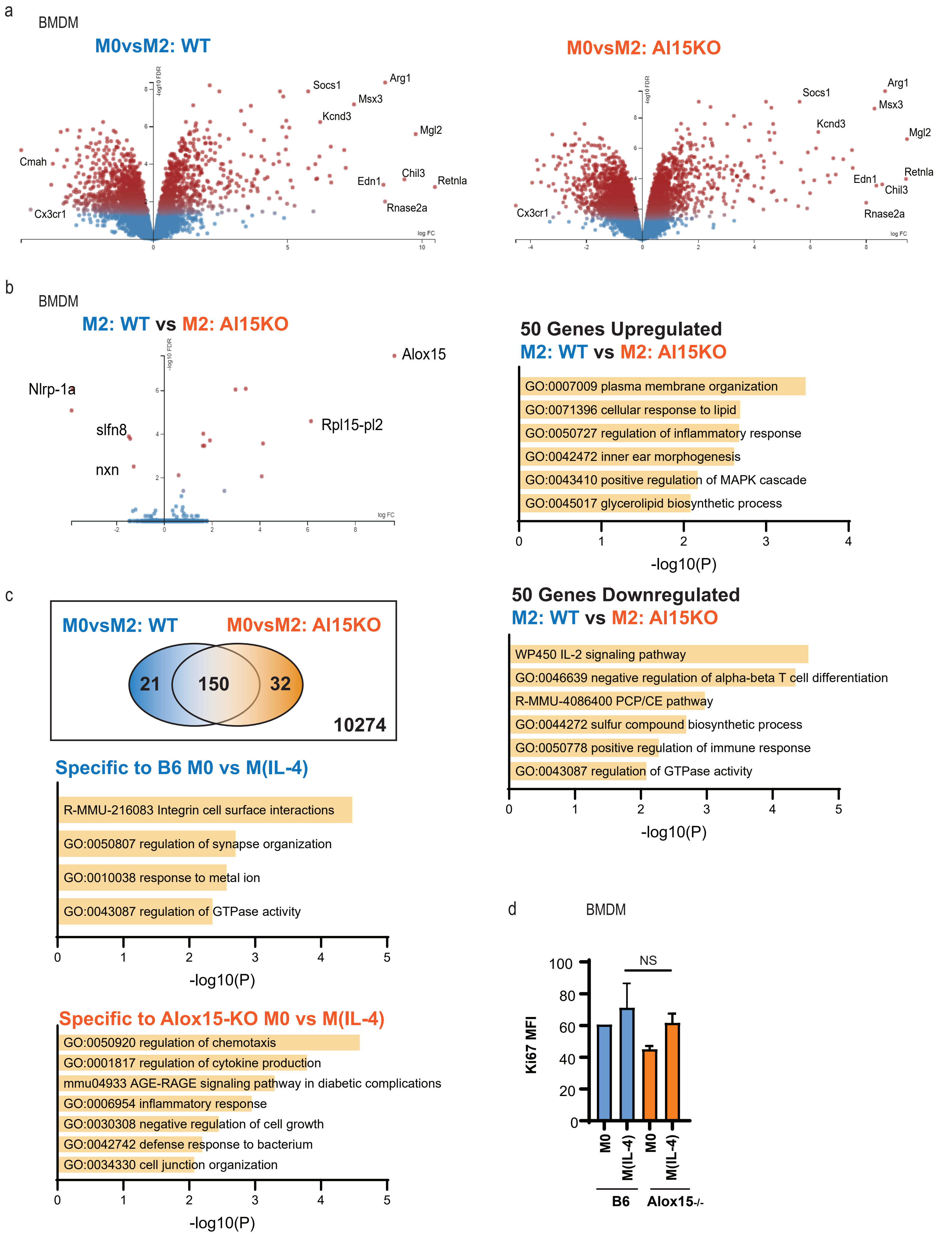
a
BMDM
M0vsM2: WT
M0vsM2: Al15KO
Socs1
Arg1
Msx3
Kcnd3
Mgl2
Cmah
Edn1
Chil3
Retnla
Rnase2a
Cx3cr1
-log10 FDR
log FC
b
BMDM
M2: WT vs M2: Al15KO
Alox15
Nlrp-1a
slfn8
Rpl15-pl2
nxn
50 Genes Upregulated
M2: WT vs M2: Al15KO
GO:0007009 plasma membrane organization
GO:0071396 cellular response to lipid
GO:0050727 regulation of inflammatory response
GO:0042472 inner ear morphogenesis
GO:0043410 positive regulation of MAPK cascade
GO:0045017 glycerolipid biosynthetic process
-log10(P)
c
M0vsM2: WT
M0vsM2: Al15KO
21
150
32
10274
50 Genes Downregulated
M2: WT vs M2: Al15KO
WP450 IL-2 signaling pathway
GO:0046639 negative regulation of alpha-beta T cell differentiation
R-MMU-4086400 PCP/CE pathway
GO:0044272 sulfur compound biosynthetic process
GO:0050778 positive regulation of immune response
GO:0043087 regulation of GTPase activity
-log10(P)
Specific to B6 M0 vs M(IL-4)
R-MMU-216083 Integrin cell surface interactions
GO:0050807 regulation of synapse organization
GO:0010038 response to metal ion
GO:0043087 regulation of GTPase activity
-log10(P)
Specific to Alox15-KO M0 vs M(IL-4)
GO:0050920 regulation of chemotaxis
GO:0001817 regulation of cytokine production
mmu04933 AGE-RAGE signaling pathway in diabetic complications
GO:0006954 inflammatory response
GO:0030308 negative regulation of cell growth
GO:0042742 defense response to bacterium
GO:0034330 cell junction organization
-log10(P)
d
BMDM
NS
Ki67 MFI
M0
M(IL-4)
M0
M(IL-4)
B6
Alox15-/-

Figure S2

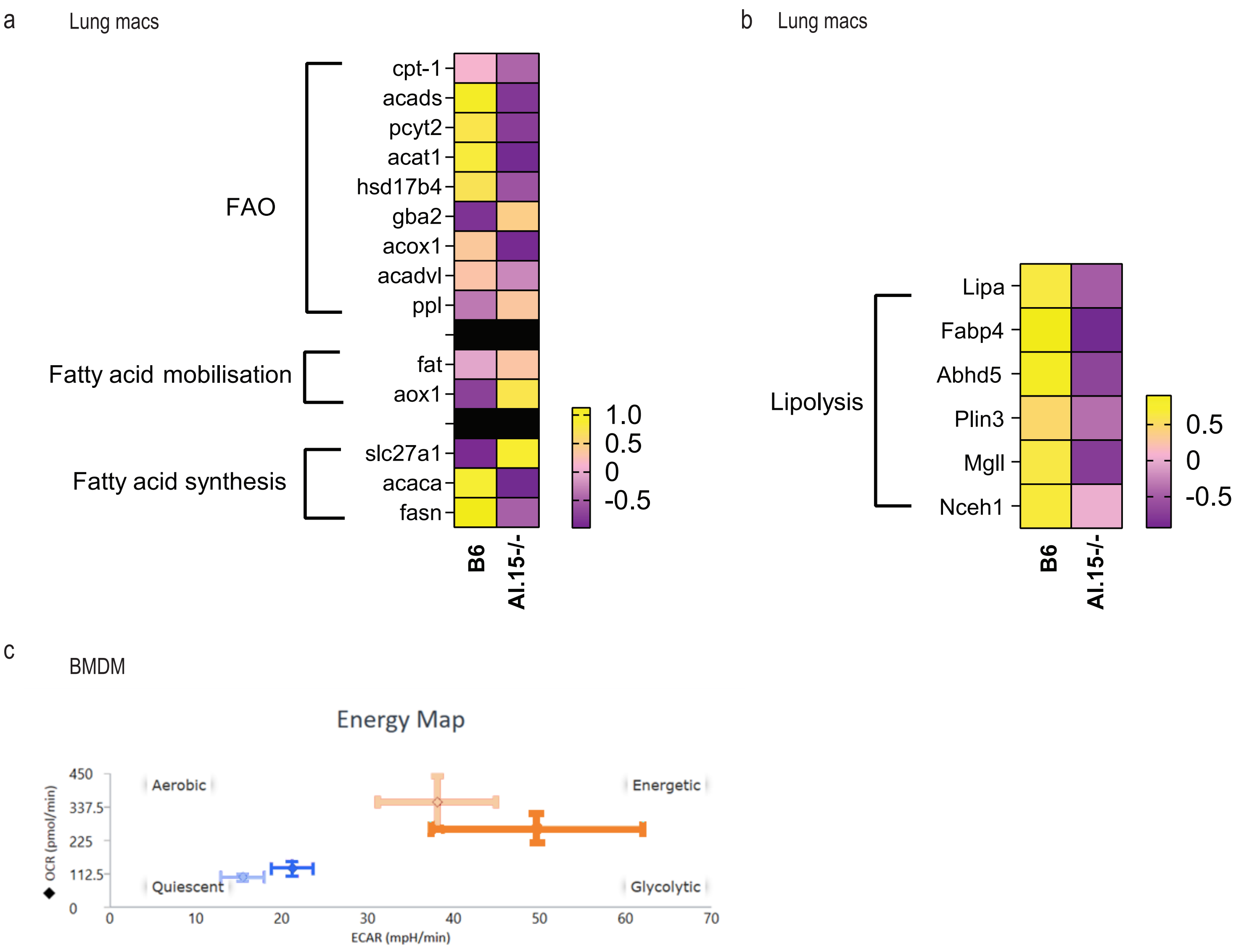
a
Lung macs
FAO
cpt-1
acads
pcyt2
acat1
hsd17b4
gba2
acox1
acadvl
ppl
Fatty acid mobilisation
fat
aox1
Fatty acid synthesis
slc27a1
acaca
fasn
1.0
0.5
0
-0.5
B6
AI.15-/-
b
Lung macs
Lipolysis
Lipa
Fabp4
Abhd5
Plin3
Mgll
Nceh1
0.5
0
-0.5
B6
AI.15-/-
c
BMDM
Energy Map
OCR (pmol/min)
450
337.5
225
112.5
0
Aerobic
Energetic
Quiescent
Glycolytic
0
10
20
30
40
50
60
70
ECAR (mpH/min)

Figure S3

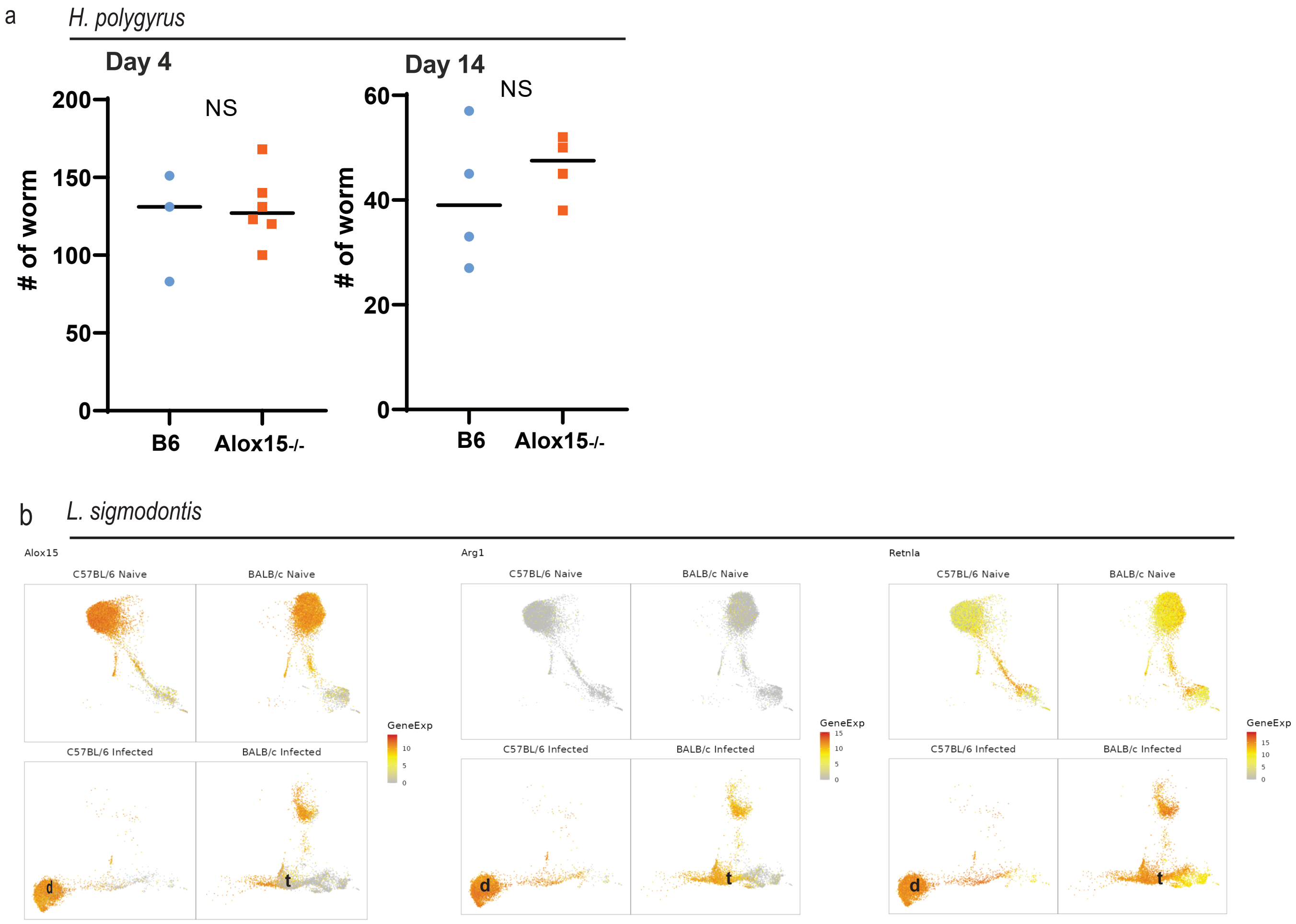
a
H. polygyrus
Day 4
Day 14
NS
NS
### of worm
200
150
100
50
0
60
40
20
B6
Alox15-/-
b
L. sigmodontis
Alox15
Arg1
Retnla
C57BL/6 Naive
BALB/c Naive
C57BL/6 Infected
BALB/c Infected
GeneExp
15
10
5
0
d
t

Figure S4

**Figure S1**

**A.** Gene expression of Alox15 was measured by qPCR in lung tissue over a 30 day timecourse after infection with 500 Nb L3 (n= 8-9 mice per timepoint, over 2 independent experiments). Macrophage specific increase of Alox15 was validated on isolated lung macrophages at D8 and D30 of Nb infection (n= 8-9 mice per timepoint, over 2 independent experiments). **B.** Survival of WT B6 mice or Alox15 deficient mice infected with a high dose of 550 L3 of Nb. N=5 per genotype. **C.** Alox15-/- and B6 controls were infected with 250 Nb L3. Two days after primary infection, larvae were collected and enumerated by baermann of lung tissue. N=4 mice per group **D.** Hemorrhage was assessed 2 days post primary infection in BALF. N=5 mice per group, pool of 2 independent experiments.

**Figure S2**

Murine bone marrow-derived macrophages were grown from B6 or Alox-15$^{-/-}$ mice and polarized or not (M0) with recombinant IL-4 (M(IL4)) for 24 hours. (n=3-4) **A.** mRNA expression profiles (volcano plots) as determined by RNA-sequencing of M0 or M(IL-4) isolated from WT B6 mice or Alox15 deficient mice. Blue, genes with not significantly differentially expressed (DE); red, genes with significant DE. FDR, false discovery rate; FC, log2 fold change. **B.** mRNA expression profiles (volcano plots) as determined by RNA-sequencing of M(IL-4) isolated from WT B6 mice or Alox15 deficient mice. Blue, genes with not significantly differentially expressed (DE); red, genes with significant DE. FDR, false discovery rate; FC, log2 fold change. Significant DE between M0 and M(IL-4) were generated using degust for both genotypes. Commonality was explored using a Venn Diagram. Pathway enrichment analysis was performed on non-common DE genes using Metascape. **C.** Flow cytometric analysis of Ki67 mean fluorescence intensity (MFI) of M0 or M(IL-4) from both genotypes. (n=3)

**Figure S3**

Total lung macrophages from B6 or Alox-15$^{-/-}$ mice 7 days post-infection with Nb L3. A. Gene expression of hallmark genes associated with fatty acid metabolic pathways (FAO, Fatty acid mobilization and Fatty acid synthesis) obtained from RNA sequencing and expressed as z-score. B Gene expression of hallmark genes associated with Lipolysis pathway obtained from

RNA sequencing and expressed as z-score. C. Energy map obtained by Seahorse flux analysis on BMDM derived from B6 or Alox-15$^{-/-}$ mice and polarized or not (M0) with recombinant IL-4 (M(IL4)) for 24 hours. (n=3-4, representative of 6 independent experiments).

**Figure S4**

**A.** Alox15-/- and B6 controls were infected with 200 Hpb L3. 4 or 14 days after primary infection, larvae were collected and enumerated by baermann of intestinal tissue. N=3-6 mice per group. B. Reanalysis of scRNA-seq of lineage-CD11b+ MNPs from naive and day 35 *L. sigmodontis*-infected resitant B6 and susceptible BALB/c mice (Finlay et al, 2023). UMAP show expression of specific genes (Alox15, Arg-1 and Retnla). The Shiny App was used for analysis (Finlay et al, 2023)

**Figure S5**

**A&C.** Murine bone marrow-derived macrophages were grown from Alox-15$^{-/-}$ mice and coculture in presence of immune serum with 100 L3 Hpb. The culture medium was or not supplemented with LOX-metabolites of arachidonic acid or increasing concentration of PPARd-g agonists. The percentage of motile larvae was assessed for each conditions by microscopy. N= 2 per condition **B.** Lungs of uninfected B6 or Alox15$^{-/-}$ mice were digested enzymatically and isolated lung Macrophages were stimulated with IL-4 with or without DPA supplementation for 24 hours. Lungs M(IL-4) were co-cultured with 100 third-stage *N. brasiliensis* larvae (Nb L3). The percentage of larvae with more than 10 cells attached was assessed for each genotypes and conditions. (n=3). **D.** SRC of BMDM stimulated as with IL-4 and/or PPARd agonist for 24 hours prior to the assay. Data representative of 3 independent experiments, n = 5 wells per conditions, each conditions with 1-3 mice per experiments.
